# KMT2D links TGF-β Signalling to Non-Canonical Activin Pathway and Regulates Pancreatic Cancer Cell Plasticity

**DOI:** 10.1101/2020.04.02.012138

**Authors:** Shuang Lu, Hong Sun Kim, Yubo Cao, Karan Bedi, Krista Chain, Lili Zhao, Ishwarya Venkata Narayanan, Zhujun Yi, Jing Yang, Yumei Gu, Michelle T. Paulsen, Mats Ljungman, Sivakumar Jeyarajan, Dafydd Thomas, Yali Dou, Howard Crawford, Marina Pasca di Magliano, Jiaqi Shi

## Abstract

Although KMT2D, also known as MLL2, is known to play an essential role in development, differentiation, and tumor suppression, its role in pancreatic cancer development is not well understood. Here, we discovered a novel signaling axis mediated by KMT2D, which links TGF-β to the activin A pathway. We found that TGF-β upregulates a microRNA, miR-147b, which in turn leads to post-transcriptional silencing of KMT2D. Loss of KMT2D induces the expression and secretion of activin A, which activates a non-canonical p38 MAPK-mediated pathway to modulate cancer cell plasticity, promote a mesenchymal phenotype, and enhance tumor invasion and metastasis in mice. We observed a decreased KMT2D expression in human primary and metastatic pancreatic cancer. Furthermore, inhibition or knockdown of activin A reversed the pro-tumoral role of KMT2D. These findings reveal a tumor-suppressive role of KMT2D and identify miR-147b and activin A as novel therapeutic targets in pancreatic cancer.

## Introduction

Pancreatic ductal adenocarcinoma (PDAC) is projected to become the second leading cause of cancer death in the United States by 2030, with an overall 5-year survival rate of less than 9%^1^. Early metastasis is one of the main reasons for poor survival for these patients. So far, no specific genetic mechanisms of PDAC metastasis have been identified. Epigenomic regulation and enhancer reprogramming are emerging as essential mechanisms for tumor progression and metastasis^3-5^. Whole-genome sequencing recently revealed frequent mutations in epigenetic regulating genes in PDAC^6,7^. Some of these mutations are considered driver mutations that lead to altered chromatin structure, promoter accessibility, and gene transcription^6,8^. However, how epigenomic dysregulation promotes PDAC progression, and metastasis is not well understood.

Whole-genome sequencing recently discovered inactivating mutations of a histone modification enzyme, KMT2D (also known as MLL2), in up to 5% of PDAC cases, suggesting its tumor suppression potential^5,9^. Inactivating mutations of KMT2D have also been associated with increased tumorigenesis and metastasis in lymphoma, esophageal, and skin cancers^9-11^, further suggesting that it is an important tumor-suppressive gene in cancer. KMT2D is one of the major histone methyltransferases for lysine 4 of histone 3 (H3K4) and mainly mono- and di-methylates H3K4 residue, which are predominant histone marks at distal promoters and enhancers^11,17^. Thus, KMT2D is known to play an essential role in establishing active promoter and enhancer landscapes, and cell-specific transcriptome. However, it remains poorly understood regarding how KMT2D contributes to PDAC progression and metastasis.

One of the critical regulators of metastasis and cancer cell plasticity is the transforming growth factor-β (TGF-β) pathway and the reversible epithelial to mesenchymal transition (EMT) process^12^. TGF-β is known to play a significant role in modulating PDAC progression^13,14^. Early in tumor development, TGF-β serves as a tumor-suppressive signal in PDAC^15-17^. Alterations in TGF-β signaling, specifically SMAD4 mutation, and inactivation, are found in more than 50% of PDAC cases. However, later in tumor progression, TGF-β induces EMT and promotes tumor invasion and metastasis^18^. The dual tumor suppressive and pro-tumorigenic role of TGF-β signaling requires a better understanding of this essential pathway during PDAC progression and metastasis.

Here, we uncovered a novel link between TGF-β signaling and epigenetic reprograming through the suppression of KMT2D expression by a microRNA, miR-147b, and the subsequent upregulation of the EMT signaling including activin, a member of the TGF-β superfamily. In addition, we have discovered that activin A promoted PDAC invasion through a non-canonical pathway via p38 mitogen-activated protein kinase (MAPK). Furthermore, inhibition of activin or p38 MAPK effectively reversed the pro-tumoral phenotype in KMT2D-knockout cells. These results not only revealed an essential novel molecular mechanism by which TGF-β promotes epigenetic reprograming, cancer cell plasticity, and PDAC progression and metastasis, but also discovered potential new therapeutic targets in PDAC patients with KMT2D inactivating mutations.

## Results

### TGF-β suppresses KMT2D expression by upregulating miR-147b

TGF-β signaling is one of the most frequently disrupted pathways in PDAC. Although steady-state transcriptional alterations induced by TGF-β have been studied, how TGF-β affects nascent RNA synthesis and stability in PDAC is not known. To better understand TGF-β signaling in PDAC, we used Bru-seq and BruChase-seq technologies to capture the changes not only in *de novo* transcriptional landscape (Bru-seq), but also RNA stability (BruChase-seq), in TGF-β treated PDAC cells^19^. As expected, genes in the TGF-β and EMT pathways were transcriptionally upregulated with the treatment (Supplementary Fig. 1a). Other upregulated pathways included hypoxia, KRAS, TNFα, UV response, unfolded protein response, and glycolysis. We also observed that TGF-β treatment downregulated transcription of genes involved in interferon-alpha, interferon-gamma, E2F, oxidative phosphorylation, G2M checkpoint, fatty acid metabolism, estrogen response, reactive oxygen species, and DNA repair pathways (Supplementary Fig. 1a). Notably, Bru-seq analysis showed that the nascent RNA synthesis of KMT2D did not change (Fig. 1a, top panel). However, BruChase-seq revealed that the KMT2D mRNA level was decreased after 6 hours of uridine chase in cells treated with TGF-β compared to control (Fig. 1a, lower panel), indicating that TGF-β treatment resulted in degradation of KMT2D mRNA. To confirm this finding, we measured KMT2D mRNA and protein levels after TGF-β treatment. TGF-β treatment reduced KMT2D mRNA (Supplementary Fig. 1b) and protein levels (Fig. 1b) in multiple PDAC cell lines. Furthermore, selective TGF-β type I receptor inhibitor, SB505124, reversed TGF-β-induced KMT2D mRNA and protein reduction (Fig. 1c and Supplementary Fig. 1b, c), confirming that TGF-β signaling inhibits KMT2D expression.

**Fig. 1.**
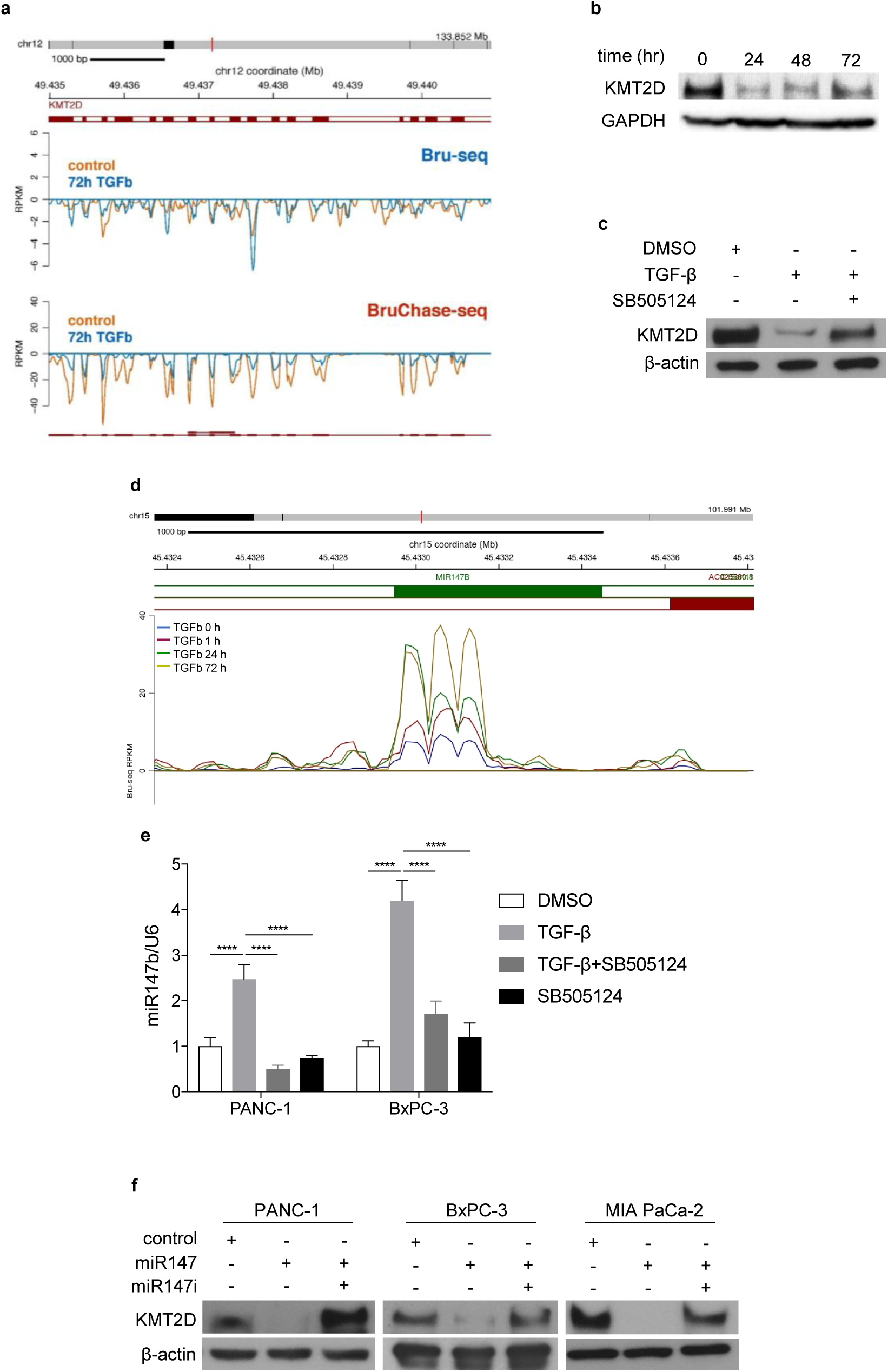
TGF-β decreases KMT2D expression through the upregulation of miR-147b. **a** Comparing KMT2D nascent mRNA synthesis (Bru-seq) and mRNA degradation (BruChase-seq) between vehicle control- and 72h TGF-β-treated PANC-1 cells. **b** Western blot analysis of KMT2D protein in PANC-1 cells treated with TGF-β for 24, 48, and 72 hours compared to vehicle-treated cells. GAPDH was used as loading control. **c** Western blot analysis of KMT2D in BxPC-3 cells treated with TGF-β with or without SB505124 for 72 hours. β-actin was used as loading control. **d** Bru-seq analysis of miR-147b transcription in PANC-1 cells treated with TGF-β for 1, 24, and 72 hours compared to vehicle-treated cells. **e** Quantitative real-time PCR of mature miR-147b in PANC-1 and BxPC-3 cells treated with DMSO or TGF-β for 48 hours with or without SB505124. U6 was used as the reference gene. (****p<0.0001, Two-way ANOVA with Dunnett’s multiple comparisons test, n=3) **f** Western Blot analysis of KMT2D in PANC-1, BxPC-3, and MIA PaCa-2 cells transfected with miR-147b mimic or miR-147b inhibitor (miR147i). β-actin was used as loading control.

Since the RNA stability of KMT2D was affected by TGF-β, we postulated that TGF-β induced microRNA degrades KMT2D mRNA. Using TargetScanHuman (www.targetscan.org), we identified microRNA-147b (miR-147b) as a candidate to target the 3’ UTR of KMT2D mRNA (Supplementary Fig. 1d). microRNA family conservation analysis showed that miR-147b is broadly conserved among mammals (Supplementary Fig. 1e), suggesting its vital function. Bru-seq data suggested that TGF-β induced miR-147b synthesis in a time-dependent manner (Fig. 1d). We hypothesized that TGF-β induces miR-147b expression, which in turn targets KMT2D mRNA for degradation. To confirm our hypothesis, we treated both PANC-1 and BxPC-3 cells with either TGF-β alone or together with TGF-β receptor inhibitor SB505124 and performed real-time PCR on steady-state RNA. TGF-β upregulated miR-147b in both cell lines, and SB505124 blocked this increase, confirming that TGF-β induces miR-147b expression (Fig. 1e). To confirm miR-147b reduces KMT2D expression, we transfected PDAC cell lines with either control miRNA, miR-147b mimic alone, or together with miR-147b inhibitor. miR-147b, but not control miRNA, inhibited KMT2D expression in all cell lines, while miR-147b inhibitor reversed this effect (Fig. 1f), supporting that miR-147b suppresses KMT2D protein expression. These results confirmed that TGF-β suppresses KMT2D expression via upregulating miR-147b.

### Genetic and protein expression alterations of KMT2D in PDAC

To assess the genetic alterations of KMT2D in PDAC, we queried The Cancer Genome Atlas (TCGA), Queensland Centre for Medical Genomics (QCMG), and UT Southwestern (UTSW) sequencing databases. We found that 5-6% of PDAC genomes carried mutations, deletions, and alterations in the KMT2D gene (Fig. 2a). Query of RNAseq data in 1,457 human cancer cell lines from The Cancer Cell Line Encyclopedia revealed significant lower expression of KMT2D mRNA in PDAC cell lines compared to all human tumor-derived cell lines (Fig. 2b)^20^. Loss of function mutations and deletions of the KMT2D gene was observed in multiple PDAC cell lines (Fig. 2c). Furthermore, patients with altered KMT2D exhibited shorter disease-free survival compared to patients with wild-type KMT2D (Fig. 2d). These data suggest that KMT2D alterations are critical in PDAC progression. To investigate the relationship between genetic alterations of the KMT2D and the TGF-β pathway, we queried the TCGA and UTSW databases. KMT2D inactivating mutations and genetic alterations of critical genes in TGF-β signaling, including TGFB1, TGFBR2, and SMAD4, are mutually exclusive in PDAC patients (Fig. 2e). These data highly suggest that the function of KMT2D and the TGF-β pathway are likely epistatic, and alteration of one gene is sufficient to drive PDAC progression.

**Fig. 2.**
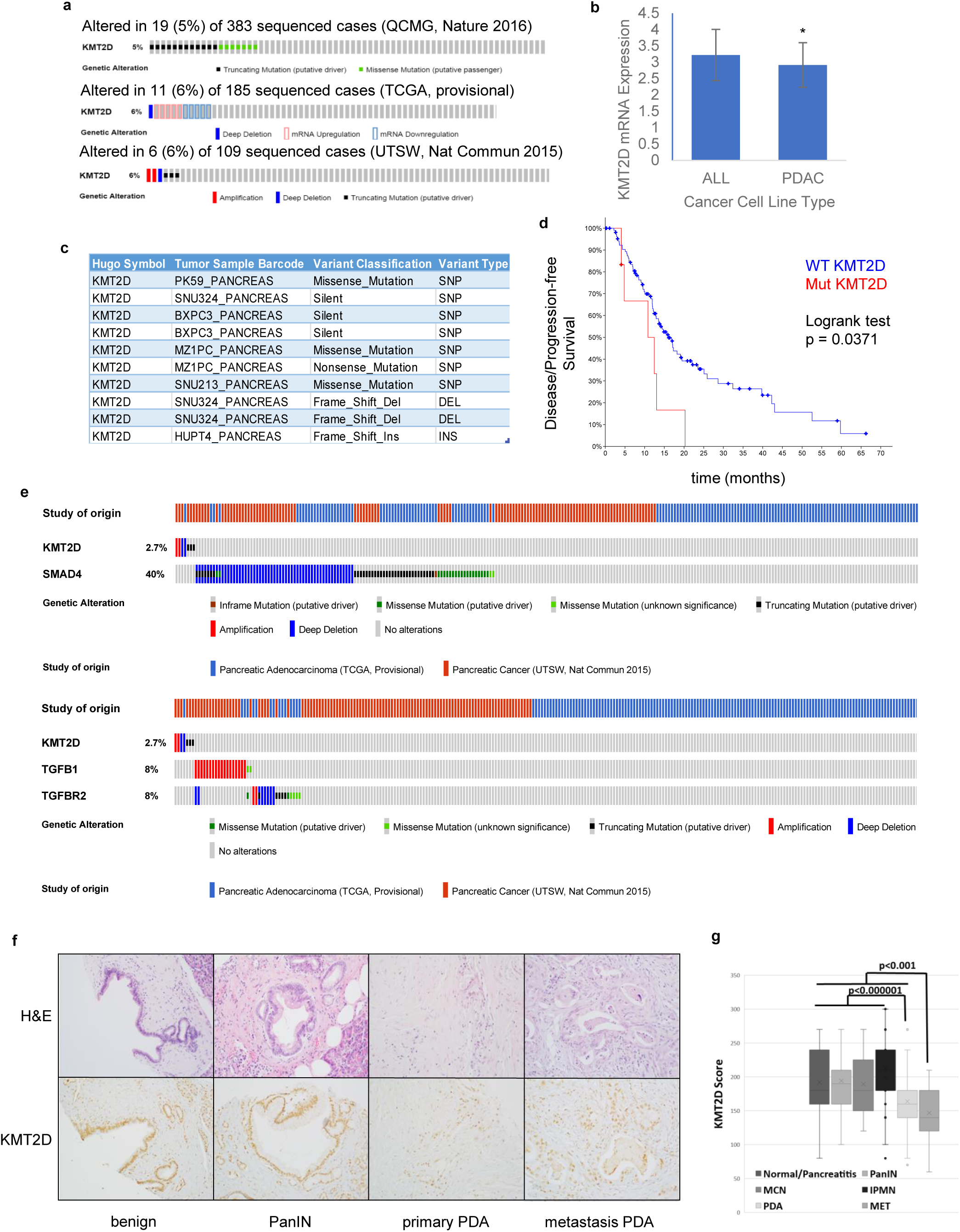
KMT2D mutation and expression profiles in PDAC patients. **a** Sequencing data of human PDACs (n=677) were queried from QCMG (top), TCGA (medium), and UTSW (bottom) databases and patients with KMT2D alterations were highlighted by various colored bars. **b** Comparison of KMT2D mRNA expression in 41 PDAC cell lines to all 1,457 human cancer cell lines based on the RNAseq data queried from the Cancer Cell Line Encyclopedia. * p<0.01. **c** Specific KMT2D mutations in representative PDAC cell lines. **d** Kaplan-Meier curve of disease-free survival in patients with either KMT2D gene mutation (Mut) or wild-type KMT2D (WT) queried from the TCGA database (n=149). **e** Patients with KMT2D, SMAD4, TGFB1, and TGFBR2 genetic alterations were queried from TCGA and UTSW databases. **f** Representative H&E and KMT2D immunohistochemical (IHC) staining in various stages of PDAC progression in tissue microarray (TMA). **g** IHC staining score of KMT2D in normal/pancreatitis (n=37), pancreatic intraepithelial neoplasm (PanIN, n=22), mucinous cystic neoplasm (MCN, n=9), intraductal papillary mucinous neoplasm (IPMN, n=36), primary (PDA, n=71) and metastatic (MET, n=14) PDAC from human TMA. Student’s t-test was used to compare between groups.

To determine whether KMT2D protein expression is also altered in human PDAC and its precursor lesions, we used a tissue microarray (TMA) containing 213 duplicated human pancreatic tissue cores of benign pancreas (n=35), precursor lesions (pancreatic intraepithelial neoplasia [PanIN], intraductal papillary mucinous neoplasm [IPMN], and mucinous cystic neoplasm [MCN]) (n=81), and primary and metastatic PDAC (n=97) (Table 1). Immunohistochemical (IHC) staining showed that KMT2D expression was lower in metastatic PDAC compared to all other lesions or benign pancreas including primary PDAC (p=0.0004), and in primary PDAC compared to all benign and precursor lesions (p=0.0000002) (Table 1 and Fig. 2f, g). These results support that KMT2D may play an essential role in PDAC progression and metastasis.

**Table 1.**
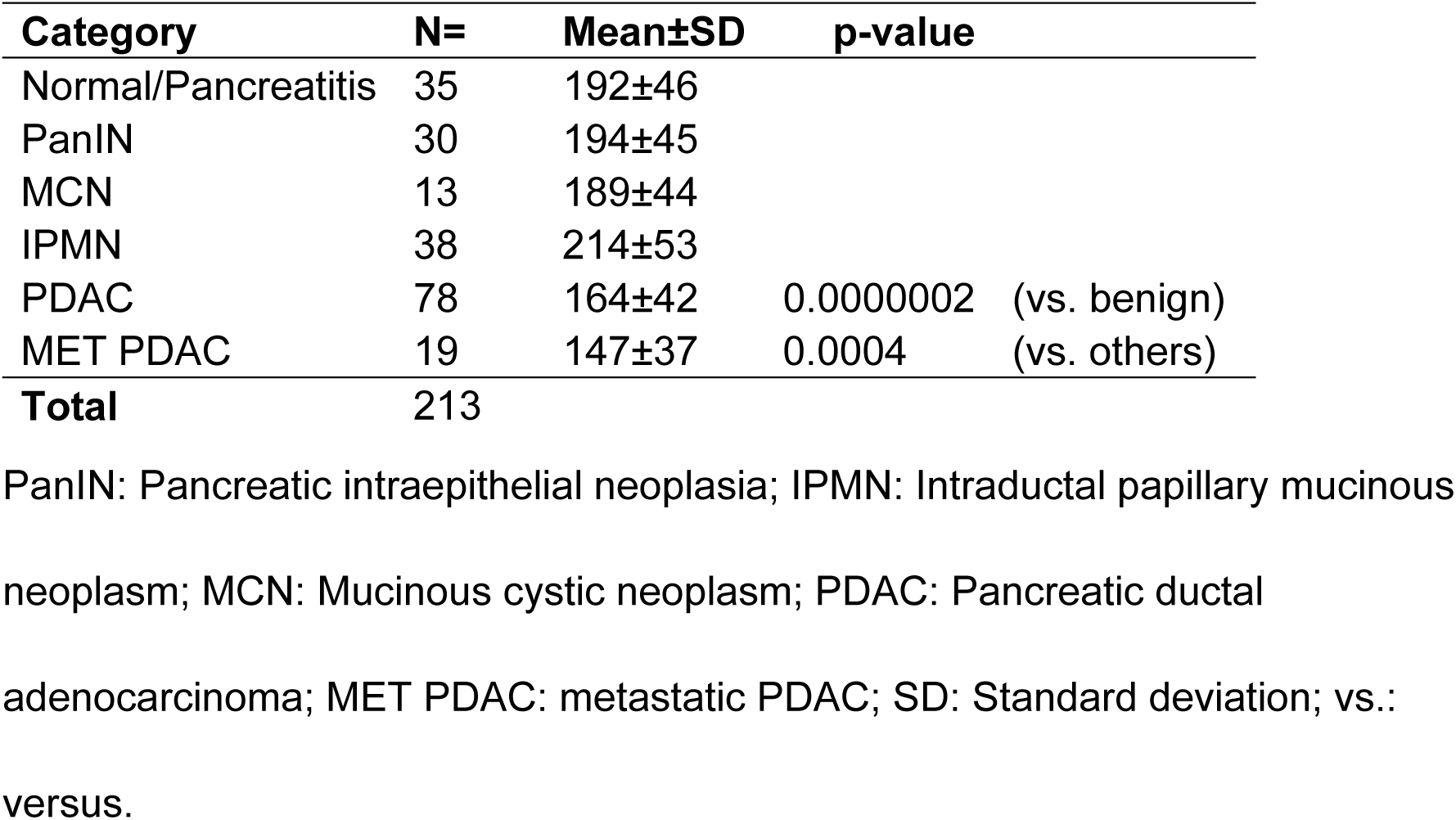
Protein expression levels of KMT2D in PDAC and its precursors.

| **Category** | **N=** | **Mean±SD** | **p-value** |
| --- | --- | --- | --- |
| Normal/Pancreatitis | 35 | 192±46 |  |
| PanIN | 30 | 194±45 |  |
| MCN | 13 | 189±44 |  |
| IPMN | 38 | 214±53 |  |
| PDAC | 78 | 164±42 | 0.0000002 (vs. benign) |
| MET PDAC | 19 | 147±37 | 0.0004 (vs. others) |
| **Total** | 213 |  |  |

PanIN: Pancreatic intraepithelial neoplasia; IPMN: Intraductal papillary mucinous
neoplasm; MCN: Mucinous cystic neoplasm; PDAC: Pancreatic ductal
adenocarcinoma; MET PDAC: metastatic PDAC; SD: Standard deviation; vs.:
versus.

### Loss of KMT2D induces a basal-like and EMT gene signature

To investigate the biologic function of KMT2D in PDAC development, we used the CRISPR/Cas9 system to generate stable KMT2D knockout PDAC cells. We then performed Bru-seq analysis to investigate the global de novo transcriptional changes in KMT2D knockout cells compared to control cells. We identified 748 common differentially transcribed genes in both knockout clones (fold change >2 or <0.5); among which, 262 genes were upregulated, and 486 genes were downregulated (Fig. 3a). Gene Set Enrichment Analysis (GSEA) showed the top pathways that were upregulated included some of the most important oncogenic and metastasis-promoting pathways like Myc, EMT, E2F, and angiogenesis (Fig. 3b). The top downregulated pathways include estrogen response, p53, cholesterol homeostasis, KRAS, and bile acid metabolism (Fig. 3c). Recently, PDAC was classified into two major molecular subtypes: classical and basal-like^21^. We next investigated whether KMT2D loss results in molecular subtype changes. We found that KMT2D knockout cells took on a more basal-like gene signature compared to control cells (Fig. 3d), which is often associated with poor prognosis. Since EMT is one of the major processes associated with TGF-β signaling, cancer cell plasticity, and metastasis, we decided to focus our study on the link between KMT2D and EMT. There was a significant enrichment of EMT signaling in both KMT2D knockout clones (Fig. 3e). Our Bru-seq analysis showed that the epithelial marker, CDH1 (encoding E-cadherin), was markedly downregulated with KMT2D loss, a hallmark of EMT (Fig. 3f top). In contrast, a mesenchymal marker, VIM (encoding Vimentin), was markedly upregulated with KMT2D loss (Fig. 3f bottom).

**Fig. 3.**
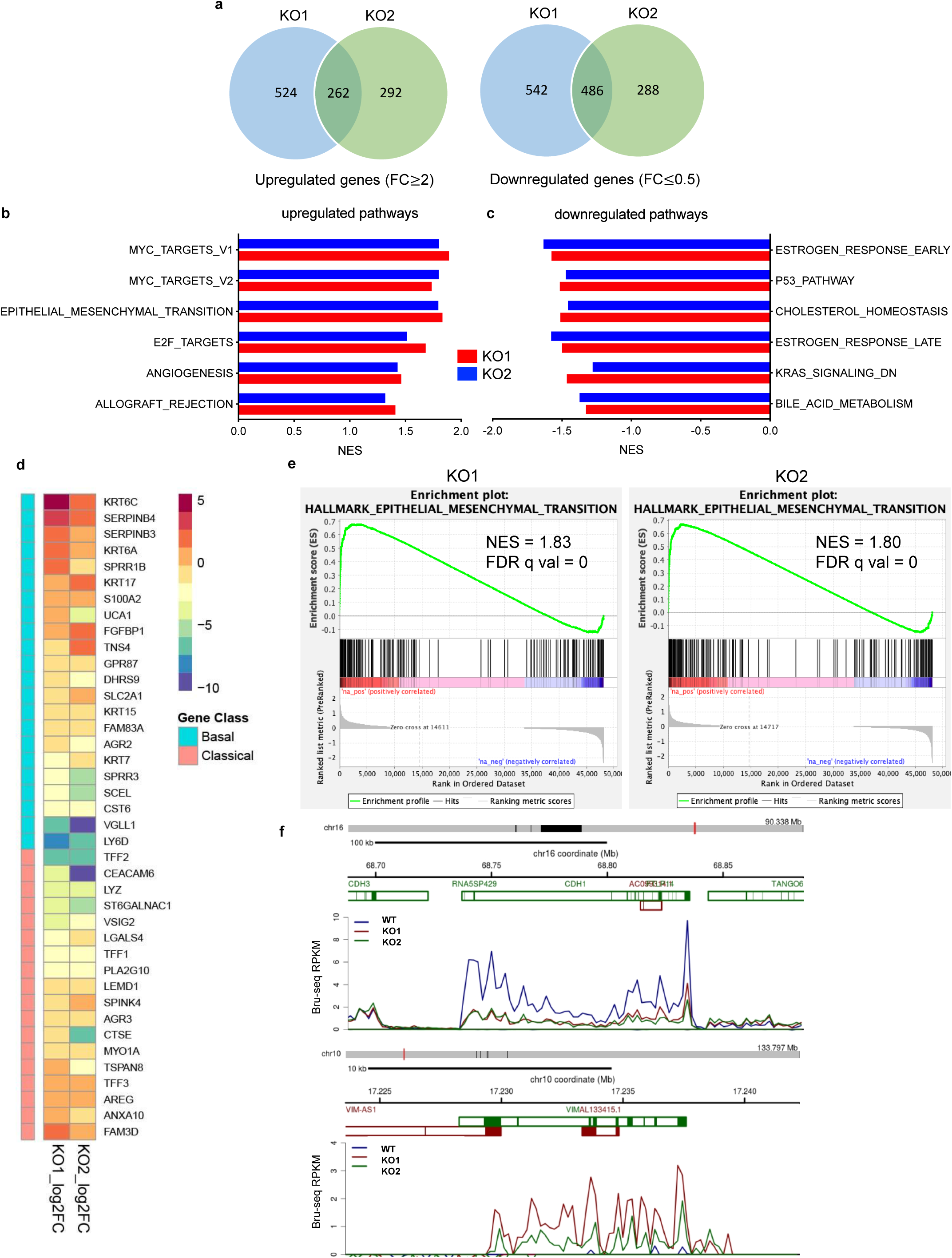
KMT2D knockout induced epigenetic reprograming. **a** Venn diagrams of differentially transcribed genes compared to control in 2 KMT2D knockout (KO) BxPC-3 clones using Bru-seq. **b, c** Top six hallmark pathways that are upregulated (**b**) or downregulated (**c**) in KMT2D KO BxPC-3 cells compared to control cells identified by Gene Set Enrichment Analysis (GSEA). **d** Heat map of altered gene signature in 2 KMT2D KO BxPC-3 clones related to the classical or basal-like subtypes of PDAC. **e** EMT pathway signature enrichment plots in 2 KMT2D KO BxPC-3 clones. NES: normalized enrichment score, FDR q val: false discovery rate. **f** Bru-seq analysis showing de novo transcripts of CDH1 and VIM in wild-type (WT) and KMT2D KO BxPC-3 cells.

To further determine the impact of KMT2D loss on PDAC cell behavior, we analyzed the changes in cell morphology, migration, invasion, and tumorigenic features. KMT2D loss induced a spindled mesenchymal cell phenotype compared to the more epithelial cell morphology in control cells (Fig. 4a), similar to the cell morphology changes induced by TGF-β treatment which was reversed by TGF-β inhibitor SB505124 (Supplementary Fig. 1f). We observed a similar change in cell morphology in PANC-1 cells with CRISPR/Cas9 knockout of KMT2D and BxPC-3 cells siRNA knockdown of KMT2D (Supplementary Fig. 2a, b). Additionally, the epithelial marker, E-cadherin, was downregulated, and mesenchymal markers, Vimentin and Snail, were upregulated in KMT2D-knockout cells compared to control cells (Fig. 4b, Supplementary Fig. 2c), indicative of EMT. The downregulation of E-cadherin and upregulation of mesenchymal markers resemble what was seen in cells treated with TGF-β and was reversible with TGF-β inhibitor SB505124 (Supplementary Fig. 1g). To confirm that these findings are due to the loss of KMT2D, we used two different siRNAs to knock-down KMT2D in 2 additional cell lines, including a patient-derived primary PDAC cell line, UM28^22^. Indeed, in both cell lines, E-cadherin expression was attenuated in KMT2D-knockdown cells (Fig. 4c), confirming the pro-mesenchymal effect of KMT2D-deficiency. We observed a global decrease in H3K4me1 and H3K4me2, but not H3K4me3, in both KMT2D knockout and siRNA knockdown cell lines (Supplementary Fig. 2d, e), which is consistent with the known function of KMT2D mainly as mono- and di-methyltransferase of H3K4.

**Fig. 4.**
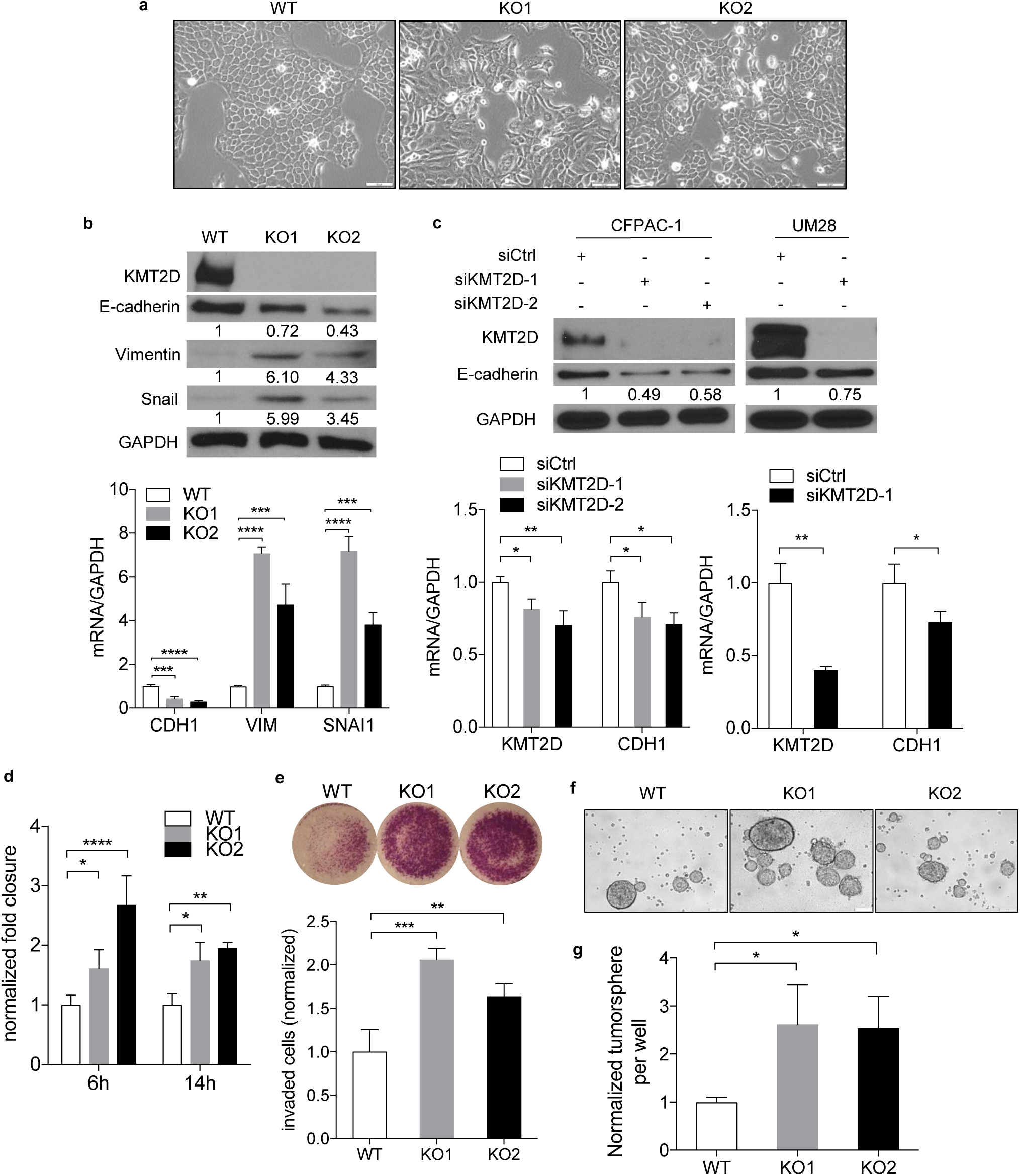
Loss of KMT2D promotes EMT, migration, invasion, and tumorigenicity. **a** Phase-contrast images of BxPC-3 wild-type (WT) and 2 KMT2D knockout (KO) clones in culture. Scale bar = 100 µm. **b** Western blot and quantitative real-time PCR of KMT2D, E-cadherin (CDH1), Vimentin (VIM), and Snail (SNAI1) expression in WT and 2 KMT2D KO BxPC-3 clones. Quantification of bands was shown relative to WT. GAPDH was used as control and reference. (***p<0.005, ****p<0.001, one-way ANOVA test with Dunnett’s multiple comparisons test, n=3) **c** Immunoblot and quantitative real-time PCR of KMT2D and E-cadherin (CDH1) in CFPAC1 and UM28 PDAC cells treated with scramble (siCtrl) or KMT2D siRNA (siKMT2D) for six days. GAPDH was used as control. Quantification of immunoblot bands were shown relative to siCtrl. (*p<0.05, **p<0.01, one-way ANOVA test with Dunnett’s multiple comparisons test (left) and unpaired student t-test (right), n=3) **d** Normalized fold wound closure in WT and KMT2D KO BxPC-3 cells at 6 and 14 hours after scratch. (*p<0.05, **p<0.01, ***p<0.005, ****p<0.001, two-way ANOVA test with Dunnett’s multiple comparisons test, n=3) **e** Invaded cells in KMT2D KO BxPC-3 cells normalized to WT at 48 hours after seeding by transwell invasion assay. (**p<0.01, ***p<0.005, one-way ANOVA test with Dunnett’s multiple comparisons test, n=3) **f** Representative pictures of WT and KMT2D KO BxPC-3 cell tumorspheres. **g** Quantification of tumorspheres in WT and KMT2D KO BxPC-3 cells. (*p<0.05, one-way ANOVA test with Dunnett’s multiple comparisons test, n=3)

Since EMT is associated with increased cell migration and invasion, we postulated that loss of KMT2D would promote cell migration and invasion. As predicted, both KMT2D knockout and knockdown cells exhibited enhanced ability of cell migration and invasion compared to control cells (Fig. 4d-e and Supplementary Fig. 2f-h). Since cancer cell plasticity is also associated with enhanced tumor-forming ability, we hypothesize that loss of KMT2D also promotes anchorage-independent cell growth potential. Indeed, KMT2D depletion led to more tumorsphere formation compared to control cells (Fig. 4f-g, Supplementary Fig. 2i). These data indicated that loss of KMT2D promotes cancer cell plasticity, cell migration, invasion, and enhanced tumorigenic potential.

### KMT2D deficiency activates activin signaling via a non-canonical pathway

To better understand how KMT2D depletion induces cancer cell plasticity and invasion, we turned our attention to the genes most consistently upregulated in KMT2D knockout cells according to our Bru-seq data. Among those genes, we found upregulated nascent INHBA transcription in both KMT2D knockout clones in comparison to an almost negligible level in control cells (Fig. 5a). Quantitative real-time PCR on steady-state RNA confirmed a roughly 3-5 fold of increase in the INHBA mRNA level in KMT2D knockout cells (Fig. 5b). Furthermore, enzyme-linked immunosorbent assay (ELISA) demonstrated an approximately higher than 9-fold increase in secreted activin A (encoded by INHBA) protein level in the media of KMT2D knockout cells (Fig. 5c). To exclude the off-target effect of the CRISPR/Cas9 system, we confirmed the upregulation of INHBA in 4 additional PDAC cell lines using KMT2D siRNA knockdown (Fig. 5d). To determine whether KMT2D deficiency induces INHBA expression by activating its promoter, we performed chromatin immunoprecipitation (ChIP)-qPCR to identify the changes of histone modification profile at the INHBA promoter region. Interestingly, despite a globally decreased H3K4me1 and H3K4me2 levels, enriched binding of H3K4me1, H3K4me2, and H3K27ac, but not H3K4me3, were observed at the INHBA promoter region in KMT2D knockout cells (Fig. 5e, Supplementary Fig. 3a). Since H3K4me1 and H3K27ac are markers of active enhancers, our results suggest that KMT2D deficiency activates INHBA expression predominantly by activating its enhancer activity.

**Fig. 5.**
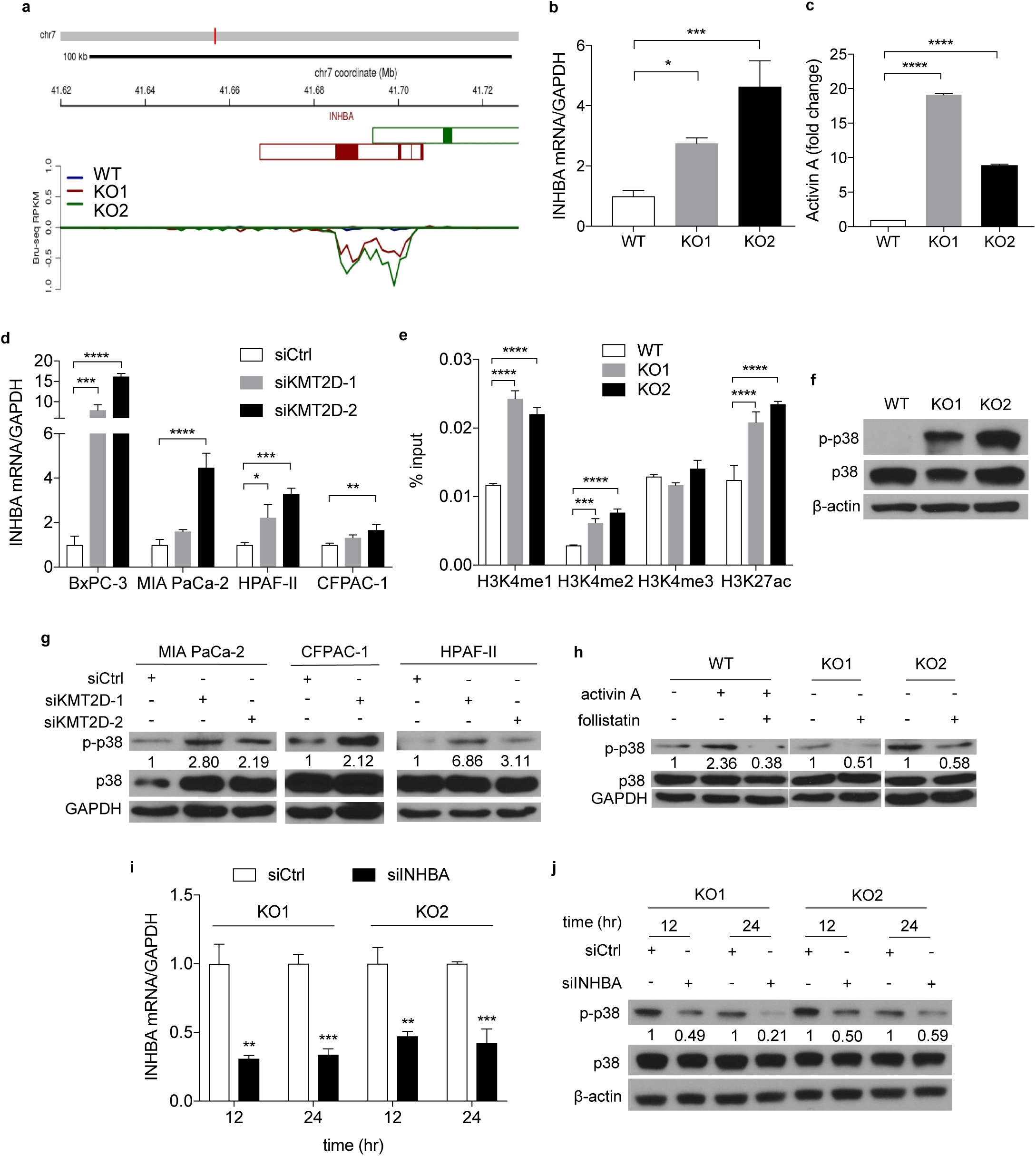
Depletion of KMT2D activates p38 MAPK by upregulating INHBA. **a** Bru-seq data of INHBA de novo transcription in wild-type (WT) and KMT2D knockout (KO) BxPC-3 cells. **b** Quantitative real-time PCR of INHBA in WT and KMT2D KO BxPC-3 cells. GADPH was used as the reference gene. (*p<0.05, ***p<0.005, one-way ANOVA test with Dunnett’s multiple comparisons test, n=3) **c** ELISA of secreted activin A protein in WT and KMT2D KO BxPC-3 cell media. (****p<0.001, two-way ANOVA test with Dunnett’s multiple comparisons test, n=2) **d** Quantitative real-time PCR of INHBA in 4 PDAC cell lines transfected with scramble (siCtrl) or 2 different KMT2D siRNAs (siKMT2D). GAPDH was used as the reference gene. (*p<0.05, **p<0.01, ***p<0.005, ****p<0.001, two-way ANOVA test with Dunnett’s multiple comparisons test, n=3) **e** Chromatin immunoprecipitation (ChIP) and real-time PCR assay of INHBA promoter region in WT and KMT2D KO BxPC-3 cells using H3K4me1, H3K4me2, H3K4me3, and H3K27ac antibodies. Results were normalized to inputs and expressed as % input. (***p<0.005, ****p<0.001, two-way ANOVA test with Dunnett’s multiple comparisons test, n=3) **f** Immunoblot of p38 and p-p38 in WT and KMT2D KO BxPC-3 cells. β-actin was used as loading control. **g** Immunoblot of p38 and p-p38 in three siCtrl or two different siKMT2D-transfected PDAC cell lines. GADPH was used as loading control. Quantification of the p-p38 band was shown relative to siCtrl. **h** Western blot analysis of p38 and p-p38 in WT and KMT2D KO BxPC-3 cells treated with or without activin A or follistatin. Quantification of the p-p38 band was shown. **i** Real-time RT-PCR of INHBA in siCtrl or INHBA siRNA (siINHBA) transfected KMT2D KO BxPC-3 cells. GADPH was used as the reference gene. (**p<0.01, ***p<0.005, unpaired student t-test, n=3) **j** Immunoblot of p38 and p-p38 in KMT2D KO BxPC3 cells transfected with siINHBA or siCtrl for 12 or 24 hours. β-actin was used as loading control. Quantification of the p-p38 band was shown relative to siCtrl.

Activin A belongs to the TGF-β superfamily and is best studied for its role in mesoderm cell fate determination during embryogenesis^23^. Similar to TGF-β, the function of activin A is highly context-dependent and has been shown to display both oncogenic and tumor-suppressive roles in different cancers. Furthermore, its role in PDAC progression is unclear. We queried the TCGA database and found that patients with altered INHBA have shorter disease-free survival compared to patients with wild-type INHBA (p=0.01, Supplementary Fig. 3b). Overall, 8% of PDAC patients have INHBA alterations, and most of these alterations are higher mRNA expression or amplification (Supplementary Fig. 3c). These data suggest that INHBA alterations are critical in PDAC progression. There are two activin A signaling cascades: canonical and non-canonical. The canonical pathway is similar to the canonical TGF-β pathway, which signals through SMAD proteins^24-26^. The non-canonical pathway includes PI3K/Akt, ERK, JNK, and p38 MAPK signaling ^27,28^. To investigate which activin A pathway KMT2D deficiency-induced, we surveyed both canonical and potential non-canonical pathways. There was no consistent change of p-SMAD2/3 in KMT2D knockout cells, suggesting that the canonical pathway is unlikely to be the mechanism (Supplementary Fig. 3d). Among non-canonical pathways, p38 MAPK was consistently activated in 4 different KMT2D knockout/knockdown cell lines (Fig. 5f, g). Additionally, activin A treatment increased p38 phosphorylation, while follistatin, an activin inhibitor, prevented p38 phosphorylation in either activin-treated or untreated KMT2D knockout cells (Fig. 5h), strongly suggesting that activin A is the mediator between KMT2D loss and p38 MAPK activation. ERK, JNK, and Akt were largely unchanged or inconsistent upon loss of KMT2D (Supplementary Fig. 3e-g). These results suggested that KMT2D deficiency likely functions through the non-canonical activin A/p38 MAPK pathway. To confirm that activin A is specifically required for the activation of the p38 MAPK pathway upon KMT2D loss, we knocked down INHBA expression (siINHBA) in KMT2D knockout cells. INHBA knockdown attenuated p38 phosphorylation in KMT2D knockout cells (Fig. 5i and 5j), confirming that loss of KMT2D activates the p38 MAPK pathway via activin A.

To determine whether activin A/p38 MAPK axis mediated the EMT and increased cell migration and invasion in KMT2D deficient cells, we knocked down INHBA in KMT2D deficient cells. The epithelial marker E-cadherin was upregulated (Fig. 6a), and tumor cell invasion was attenuated in INHBA knockdown cells (Fig. 6b). Furthermore, we treated KMT2D deficient cells with p38 MAPK inhibitor SB202190 and observed the upregulation of E-cadherin (Fig. 6c) and decreased tumor cell migration and invasion in a dose-dependent manner (Fig. 6d-e). As evidence of treatment effectiveness, the phosphorylation of MAPK-activating protein kinases 2 (MK2), a well-characterized downstream target of p38, was decreased upon SB202190 treatment (Fig. 6c). In comparison, p38 MAPK inhibitor failed to upregulate E-cadherin in KMT2D preserved cells (Supplementary Fig. 3h). These results support that KMT2D depletion-induced EMT, migration, and invasion are, at least in part, mediated by activin A/p38 MAPK pathway.

**Fig. 6.**
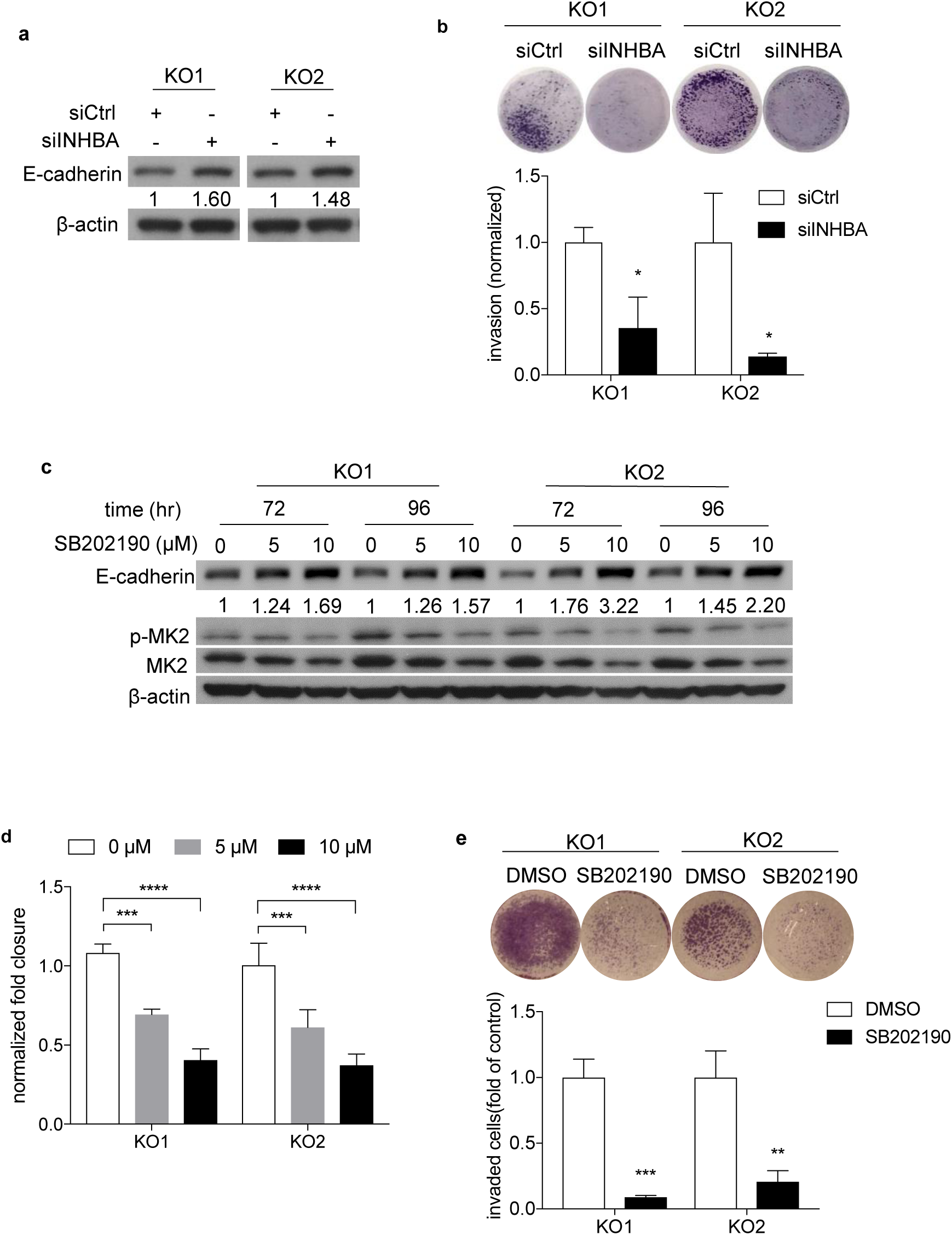
Depletion of INHBA or inhibition of p38 MAPK upregulated E-cadherin expression and attenuated PDAC cell migration and invasion. **a** Immunoblot of E-cadherin in KMT2D KO BxPC-3 cells transfected with either siCtrl or siINHBA. β-actin was used as loading control. Quantification of E-cadherin bands was shown. **b** Transwell invasion assay in KMT2D KO BxPC-3 cells transfected with siCtrl or siINHBA. (*p<0.05, unpaired student t-test, n=3) **c** Immunoblot of E-cadherin, MK2, and p-MK2 in KMT2D KO BxPC-3 cells treated with 5 or 10 µM of SB202190 or DMSO control for 72 or 96 hours. β-actin was used as loading control. Quantification of E-cadherin bands was shown. **d** Normalized fold closure in DMSO or 5 or 10 µM SB202190-treated KMT2D KO BxPC-3 cells by wound healing assay. (***p<0.005, ****p<0.001 Two-way ANOVA test with Dunnette’s multiple comparisons, n=3) **e** Transwell invasion assay of KMT2D KO BxPC-3 cells treated with either DMSO or 5 µM SB202190. (**p<0.01, ***p<0.005, unpaired student t-test, n=3)

### Loss of KMT2D promotes tumor growth, metastasis, and EMT in vivo

To validate our findings *in vivo*, we injected KMT2D knockout or wild-type PDAC cells orthotopically into the pancreas of NOD-SCID IL2Rgamma^null^ (NSG) mice. Mice implanted with KMT2D knockout cells developed bigger primary tumors (Fig. 7a, b) and metastases (Fig. 7c) compared to the control group. KMT2D knockout tumors had more mesenchymal or basal-like histology with decreased epithelial marker, E-cadherin, and increased mesenchymal marker, Vimentin, compared to control tumors (Fig. 7d-e, Supplementary Fig. 4a-b), confirming the EMT process induced by KMT2D deficiency observed *in vitro*. Furthermore, we observed that p-p38 levels increased in tumors derived from KMT2D knockout cells compared to wild-type tumors (Fig. 7f-g). In summary, our *in vivo* data supports our hypothesis that loss of KMT2D promotes metastasis and tumor progression by activating activin A/p38 MAPK pathway and EMT process.

**Fig. 7.**
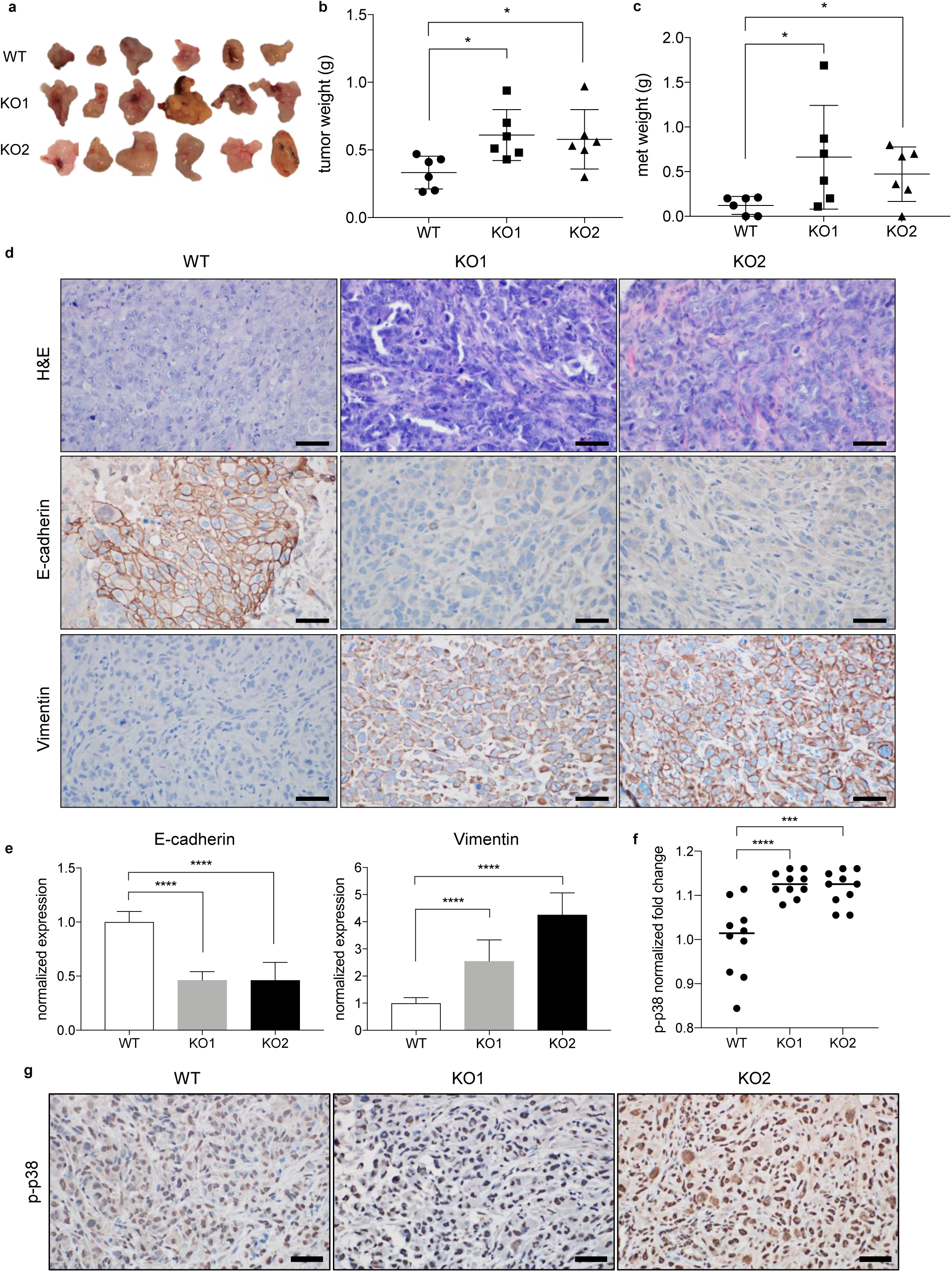
KMT2D depletion promotes both primary and metastatic PDAC growth and EMT in vivo. **a** Tumor harvested from WT and KMT2D KO BxPC-3 cell orthotopic xenograft four weeks after injection (n=6 each group). **b, c** Weight of primary (**b**) and metastatic (**c**) tumors harvested from WT and KMT2D KO orthotopic xenograft mouse model (*p<0.05, unpaired student t-test, n=6). **d** Representative H&E and IHC stains of E-cadherin and Vimentin in WT and KMT2D KO PANC-1 orthotopic xenograft tumors. (scale bar = 50 µm) **e** Quantification of IHC stains of E-cadherin and Vimentin in WT and KMT2D KO tumors. (**p<0.01, ***p<0.005, ****p<0.001, One-way ANOVA test with Tukey’s multiple comparisons, n=30). **f** Number of p-p38 positive cells in WT and KMT2D KO PANC-1 cell orthotopic xenograft tumors. Five random fields per animal were examined. Each data point represents one field. (***p<0.005, ****p<0.001, One-way ANOVA test with Dunnett’s multiple comparisons, n=10) **g** Representative pictures of p-p38 IHC staining in PANC-1 xenograft tumor.

## Discussion

Treating pancreatic cancer patients is particularly challenging due to the early occurrence of metastasis. Multiple groups performed comparative genomic analysis between matching primary and metastatic tumors but failed to find metastasis-specific gene^5,29^. This observation led to the hypothesis that epigenetic alterations may regulate PDAC metastasis. Indeed, chromatin remodeling epigenetic regulators, including KMT2D, were identified as potential drivers for pancreatic cancer progression^6^. However, our understanding on how epigenetic dysregulation affects pancreatic cancer behavior is still elementary. Here, we discovered KMT2D as a new critical link between the TGF-β and the non-canonical activin signaling to promote EMT and PDAC progression (Fig. 8).

**Fig. 8.**
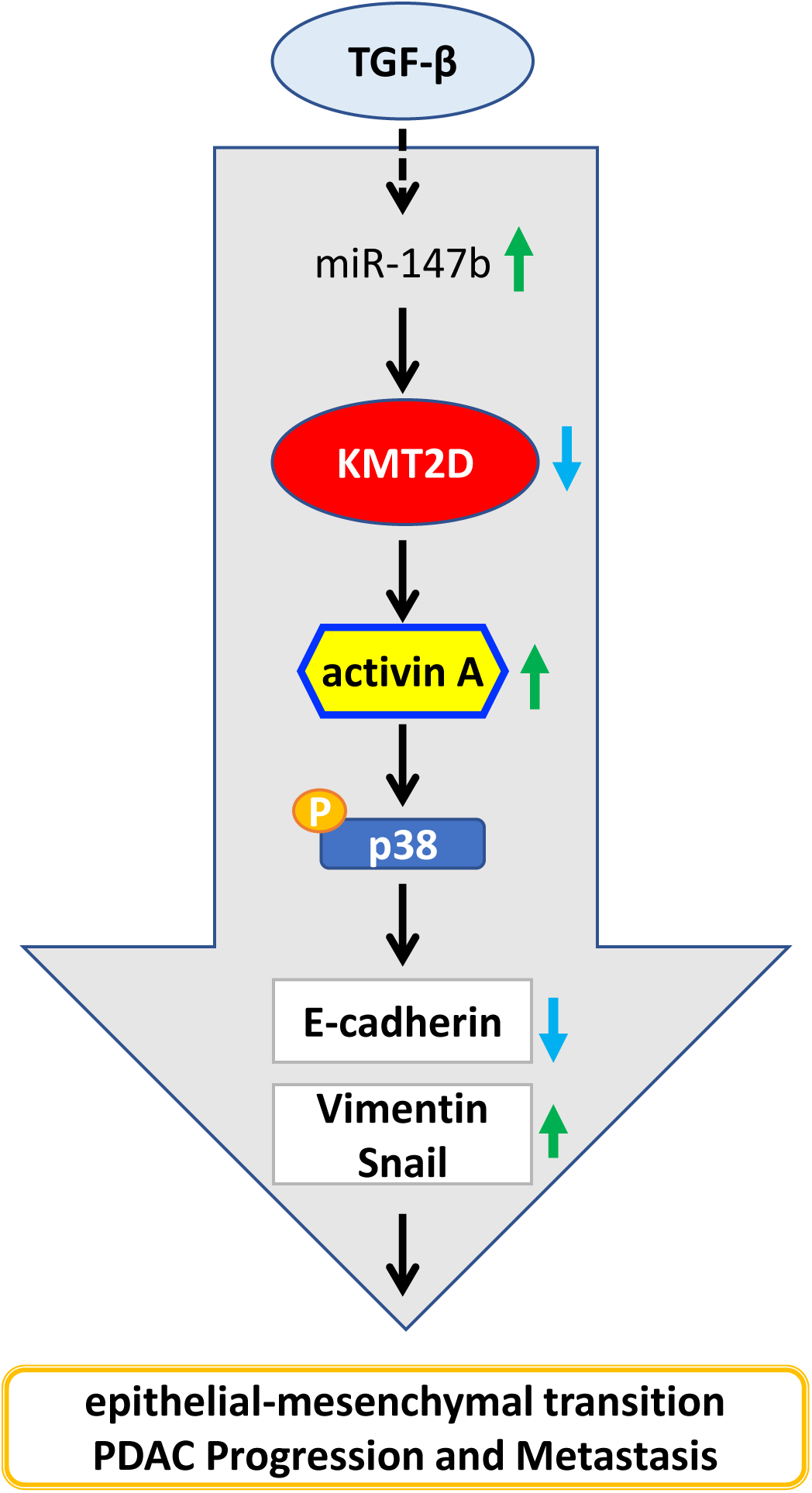
Overall hypothesis: KMT2D links TGF-β and activin A signaling. TGF-β induces KMT2D mRNA degradation by the upregulation of miR-147b. Decreased expression of KMT2D subsequently upregulates the transcription of INHBA/activin A, which then promotes EMT, cell migration and invasion, tumorigenicity, PDAC progression, and metastasis via a p38-mediated non-canonical pathway.

TGF-β is one of the most critical cytokines in pancreatic carcinogenesis. However, its highly context-dependent dual pro- and anti-tumor function is not fully understood and thus hindered the development of targeting strategies. Elevated TGF-β level is associated with poor survival in pancreatic cancer patients^30^. Tumor supportive role of TGF-β in pancreatic cancer sparked much interest in search for metastasis driving genes regulated by TGF-β. Global evaluation of the effect of TGF-β on PDAC cells has been performed on steady-state gene expression using either conventional high throughput RNA-Seq or gene microarray analysis^14,31^. However, the effect of TGF-β signaling on pancreatic cancer epigenetics is not well studied. Other groups have reported that TGF-β induces EMT through the upregulation of epigenetic regulators, KDM4B^32^ and KDM6B^33^. However, past studies did not identify how TGF-β regulated the expression of these epigenetic regulators. In this study, we used Bru-seq and BruChase-seq technologies to obtain a readout of RNA synthesis and stability in TGF-β treated cells. Bru-seq technology measures *de novo* RNA transcription independently from pre-existing RNAs (Bru-seq), as well as RNA stability (BruChase-seq)^19^. Using this unique technology, we discovered that TGF-β downregulates epigenetic regulatory protein, KMT2D, expression by post-transcriptional gene silencing.

Much work has been done to characterize distinct microRNAs expressed in pancreatic cancer compared to normal tissue^34-36^. We report here that miR-147b expression increases with TGF-β treatment, attenuates KMT2D expression, and subsequently allows the cancer cells to acquire a mesenchymal phenotype. Others have also reported the pro-tumor role of miR-147b^37-39^. For example, miR-147b increases the tumorigenicity in hepatocellular carcinoma by targeting ubiquitin-conjugating enzyme E2N^37^. In patients with hepatitis C-associated diffuse large B-cell lymphoma, high miR-147b level is correlated with poor prognosis^38^. Recent work in lung cancer showed that miR-147b endows enhanced drug resistance by altering tricarboxylic acid cycle^39^. These findings suggest the possibility of using miR-147b as a therapeutic target for multiple tumor types.

Alterations in the KMT2D gene have been reported for multiple types of cancers^40^. Inactivating KMT2D mutations have been associated with poor prognosis of some cancers such as non-small-cell lung cancer^41^, while high KMT2D expression predicts poor prognosis and promotes tumor progression in other cancers, including esophageal squamous cell carcinoma^42^. Dawkins and colleagues reported that reduced expression of KMT2D is associated with improved survival of PDAC patients^43^. However, a more recent study by Koutsioumpa and colleagues showed that low KMT2D expression increased proliferation and tumorigenicity in PDAC by altering glucose and lipid metabolism^44^. Our study showed that primary and metastatic PDACs had significantly lower expression of KMT2D and loss of KMT2D promoted PDAC invasion and migration, which supports a tumor-suppressive role of KMT2D in PDAC. Loss of KMT2D also induced a gene signature closely resembling the Moffitt basal-like subtype, which confers a significantly worse prognosis compared to the classical subtype^21,45,46^. These PDAC subtypes have distinct epigenetic landscapes that drive transcriptional alterations. Our study has suggested that KMT2D is a crucial epigenetic regulator that contributes to the basal-like subtype of PDAC. Furthermore, we discovered that activin A pathway was activated by KMT2D loss through a non-canonical signaling via p38 MAPK.

Activin A belongs to the TGF-β superfamily and is best studied for its role in mesoderm cell fate determination during embryogenesis^47,48^. However, its role in pancreatic cancer development is not well studied. Similar to TGF-β, activin A post-natal expression is tightly regulated and highly tissue-specific. Normal pancreas expresses negligible activin A, according to the Human Protein Atlas. Activins are homo- or hetero-dimers of activin β subunits. Currently, there are three known bioactive activin dimers: activins A (β_A_β_A_), B (β_B_β_B_) and AB (β_A_β_B_)^49,50^. After secreted from the cell, activin A binds to ActRII/IIB, which recruits ALK4 to phosphorylate SMAD2/3 in a similar mechanism to TGF-β^24-26^. In addition to the canonical pathway, non-canonical pathways, such as PI3K/Akt, ERK, JNK, and p38, have been associated with activin A function independent of SMAD activation^27,28^. In our study, we found that activin A and p38 MAPK are required for loss of KMT2D-induced EMT and enhanced tumor cell invasion since inhibition or knockdown of either activin or p38 MAPK attenuated EMT and tumor cell invasion induced by KMT2D loss. These findings revealed activin A and p38 MAPK as novel potential therapeutic targets for PDAC patients with low KMT2D.

In summary, we uncovered an essential crosstalk between TGF-β, KMT2D, and activin signaling to promote PDAC cell plasticity, basal-like gene signature, tumorigenicity, invasion, and metastasis. These findings shed light on the poorly understood functions of KMT2D and activin signaling in PDAC progression and filled in the gap of our knowledge on TGF-β function. More importantly, we identified miR-147b and activin A as novel potential therapeutic targets in PDAC.

## Methods

### Cell culture and transfections

Cell lines were cultured in the appropriate medium supplemented with 10% fetal bovine serum and 1% penicillin/streptomycin (GIBCO) at 37°C with 5% CO_2_. KMT2D siRNAs or scrambled siRNA (Dharmacon) (50 nM) were transfected into PDAC cells using Lipofectamine RNA iMAX reagent (Invitrogen) according to the manufacturer’s instructions. Synthesized guide RNAs (sgRNAs) and Cas9 protein (Dharmacon) were transfected into PANC-1 and BxPC-3 cell lines by RNA iMAX, followed by clonal cell isolation. Briefly, a single CRISPR/cas9-edited cell was plated in 96-well plate using limiting dilution method, and clonal cells were identified and expanded. The sequences of siRNAs and sgRNAs are listed in Supplementary Table 1.

### Reagents and antibodies

All reagents and antibodies are listed in Supplementary Table 2.

### Bru-seq and BruChase-seq

Bru-seq and BruChase-seq were performed as previously described^19^. Briefly, PANC-1 cells were incubated in media containing bromouridine (Bru) (Sigma-Aldrich) at a final concentration of 2 mM for 30 minutes at 37°C to label nascent RNA. Cells were then lysed directly in Trizol, and total RNA was isolated. Bru-labeled RNA was immunocaptured using anti-BrdU antibodies, followed by the preparation of strand-specific cDNA libraries with the Illumina TruSeq kit (Illumina) and deep sequencing using the Illumina sequencing platform as previously described^19,51,52^. For BruChase-seq, cells were first labeled with 2 mM Bru for 30 minutes, washed in PBS, and then incubated in conditioned media containing 20 mM uridine for 6 hours. The cells were then lysed in Trizol, and Bru-labeled RNA was captured and processed as described above. log2FoldChange values from Bru-seq analysis were plotted for the two knockouts (vs. control) for the basal and classical gene sets to create a heat map. The heat map was generated in R using the pheatmap package (v1.0.12).

### Protein extraction and western blot analysis

For the preparation of cellular lysates, cultured cells were harvested with NP-40 based whole-cell lysis buffer (50 mM Tris, pH 8.0, 150 mM NaCl, 2 mM EDTA, 1 mM PMSF, 1X proteinase inhibitor and 1.5% NP-40) as previously described^53^. Protein concentrations were measured using the Bradford assay (Bio-Rad). Protein samples were heated and separated on sodium dodecyl sulfate (SDS)-PAGE or Tris-Acetate-PAGE gel-electrophoresis and transferred to PVDF membranes (Millipore). After blocking in phosphate-buffered saline (PBS)/Tween-20 containing 5% nonfat milk, the membranes were incubated with primary antibodies overnight at 4°C, followed by incubation with peroxidase-conjugated secondary antibodies (Jackson ImmunoResearch Laboratories) for 1 hour. Then the proteins were visualized using an ECL detection kit (Thermo Scientific).

### RNA preparation and Quantitative Real-Time PCR

Total RNAs were obtained using the RNeasy Mini Kit (QIAGEN), and microRNAs were purified by the miRNeasy Mini Kit (QIAGEN). Reverse transcription of RNA was performed using the SuperScript ® III First-Strand Synthesis kit (Invitrogen, Carlsbad, CA) while the reverse transcription of microRNA was obtained using TaqMan™ MicroRNA Reverse Transcription Kit according to the manufacturer’s instructions. Quantitative real-time RT-PCR was used to quantify RNA and microRNA expression using SYBR Green reagents (Applied Biosystems) and microRNA with TaqMan™ Universal Master Mix II (Applied Biosystems) in MicroAmp Optical 96-well reaction plates (Applied Biosystems). The 2^-ΔΔCT^ method was used to measure the gene expression compared with the endogenous controls (U6 non-coding small nuclear RNA for miR-147b and GAPDH for mRNAs). Primers for miR-147b and U6 were purchased from Thermo Fisher Scientific, and other primers were designed using Primer-BLAST (https://www-ncbi-nlm-nih-gov.proxy.lib.umich.edu/tools/primer-blast/) and synthesized by Integrated DNA Technologies. The primer sequences are listed in Supplementary Table 3.

### Patient pancreatic tissue samples and clinicopathologic variables

The Institutional Review Board at the University of Michigan approved the study (protocol number: HUM00098128). Patients with pancreas resections for pancreatitis, cystic neoplasms, or PDAC from 2002 to 2015 at the University of Michigan Health System were included in the study. All pathology slides were reviewed, and the diagnosis confirmed by a gastrointestinal pathologist.

### Tissue microarrays

All Hematoxylin and Eosin (H&E) stained slides were reviewed, and diagnoses confirmed by a gastrointestinal pathologist (JS) and corresponding areas were carefully selected and marked. Duplicated 1 mm diameter tissue cores from a total of 311 patient tissue samples were selectively punched/extracted and transferred to recipient tissue array blocks. Five tissue microarrays (TMAs) were set up according to a standard protocol, as previously described^54^. H&E staining was performed on each TMA block using standard protocol, and unstained slides were prepared for immunohistochemical staining.

### Immunohistochemical analysis and scoring

IHC staining of the TMAs was completed using the KMT2D antibody (MilliporeSigma, Burlington, MA) and, as previously described^54^. The intensity of the staining was recorded as: 0 = negative; 1 = weak; 2 = moderate; 3 = strong. The percentage of positive cells was also assessed. The final IHC staining score was calculated as the staining intensity multiplied by the percentage of positive cells. The cases were divided into KMT2D high versus low groups with the median as the cutoff. IHC staining of E-cadherin and Vimentin was quantified using ImageJ software. Five random fields of the primary tumor were chosen for quantification. p-p38 staining was quantified by counting positive cells in 100 tumor cells in 5 randomly selected fields in the primary tumor. Each data point represents an individual field that was analyzed.

### In vitro wound-healing assay

Cell migratory ability was measured by wound healing assay, which was performed by seeding cells in complete media in 6-well plate for 24-48 hours until a confluent monolayer had formed. Linear scratches were made using a sterile 20 μL pipette tip. Monolayers were washed three times with PBS to remove detached cells, then the cells were cultured in complete media and incubated in a humidified incubator at 37°C. Photographs of the wound were taken immediately after wound formation and at the end of the assay with phase-contrast microscopy. The wound area was measured over time using ImageJ.

### In vitro transwell invasion assay

Cell invasion assay was performed using a 24-well plate with 8-μm pore size chamber inserts (Corning). 0.5 - 1×10^5^ cells in serum-free culture medium were seeded into the upper chamber per well with a Matrigel-coated membrane. 500 μL of complete growth medium was added to each lower chamber. After incubation for 48 hours at 37°C, cells that invaded through the membrane were fixed and stained with the DIFF staining kit (IMEB INC), photographed, and counted using ImageJ. Representative whole membranes were imaged under the same conditions and shown.

### Tumor sphere cultures

Single cells were suspended in X-VIVO 10 Serum-free Hematopoietic Cell Medium (#04-380Q, Lonza) in six-well Ultra-Low Attachment Plates (Corning). For the KMT2D knockout BxPC-3 cell line, single cells from low passage BxPC-3 wild type and two knockout clones were suspended and seeded into the plates for five days. For the KMT2D siRNA knockdown UM28 cell line, attached cells were transfected with the scrambled siRNA or KMT2D-targeted siRNA 24h before they were suspended and seeded into the wells for five days. Pictures of the tumorsphere were taken using an inverted microscope (Olympus) and quantified manually.

### Chromatin Immunoprecipitation-qPCR (ChIP -qPCR)

ChIP was performed using the SimpleChIP Enzymatic Chromatin IP Kit (#9003 Cell Signaling) according to the manufacturer’s instructions with minor modifications. In brief, approximately 4 × 10^6^ cells per IP were formaldehyde-crosslinked, and chromatin was digested into 100-400 bp fragments by micrococcal nuclease. Then the digested chromatin was immunoprecipitated with H3 (#9003 Cell Signaling), IgG (#9003 Cell Signaling), H3K4me1(ab8895 Abcam), H3K4me2(#17677 EMD Millipore), H3K4me3(#9751 Cell Signaling), or H3K27ac (ab4729 Abcam) antibodies using the recommended dilutions. Washed and purified DNA were subjected to qPCR using SYBR Green reagents (Applied Biosystems) according to the manufacturer’s instructions. The primers directed against the INHBA promoter region were designed and used in qPCR reactions. The enrichment of the PCR product in the target DNA fragment in each condition was compared to the % of DNA quantity in the input sample. The % input was calculated as follows: Percent Input = efficiency x 2^(C[T] 2%Input Sample - C[T] IP Sample)^.

### Establishment of orthotopic xenograft mouse model

Briefly, 6-8 weeks old NSG mice were anesthetized with isoflurane. An incision was made at the left abdominal flank, close to the splenic silhouette. The pancreas was pulled out of the peritoneal cavity and exposed for injection. 1×10^6^ BxPC-3 or PANC-1 cells were suspended in 40 μL mixed cell culture media and Matrigel (BD) at a 1:1 ratio and injected into the tail of the pancreas. The abdominal muscle and skin layers were sequentially closed with sutures. The animals were monitored every day for one week after surgery for any adverse effects, and once a week for four weeks to check for tumor growth. The study was terminated four weeks after tumor implantation surgery. Tumor tissue was harvested, fixed in formalin, and then embedded in paraffin. Subsequently, 5 μm paraffin sections were obtained and underwent H&E and IHC staining. All studies were carried out under the protocol approved by the Institutional Animal Care and Use Committee (IACUC) at the University of Michigan.

### Statistical analysis

ANOVA models were used to compare the expression of KMT2D in PDAC and its precursor lesions. Analysis was performed using GraphPad Prism software or SAS (version 9.4, SAS Institute) or Excel for all in vitro and in vivo studies. Results are presented as mean ± SD unless otherwise noted. Statistical significance among different groups was calculated as described in each figure. Significance is determined if p<0.05.

## Data Availability

All data needed for interpretation of the results are presented in this paper and its supplementary information files. Source data underlying Figs. 1c, 1d, 1f, 1g, 2b, 2g, 4b-e, 4g, 5b-i, 6a-e, 7b-d, 7f, and Supplementary Figs. 1a-b, 1f, 2c-I, 3a, 3d-k, 4b have been made available upon request. The datasets generated during and/or analyzed during the current study are available from the corresponding author on reasonable request.

## Acknowledgments

The JS lab is funded by the National Cancer Institute (K08 CA234222). The authors would like to thank the lab members of Drs. Mats Ljungman, Yali Dou, Howard Crawford, Marina Pasca di Magliano, Costas Lyssiotis, James Moon, and Timothy Frankel for sharing reagents, resources, and providing advice. We thank Dr. Kai Ge for kindly providing us the KMT2D antibody.

## Author information

These authors contributed equally: Shuang Lu, Hong Sun Kim, Yubo Cao.

## Contributions

Conceptualization: J.S.; designed the experiments: S.L., H.S.K, Y.C., M.L., and J.S.; performed the experiments: S.L., H.S.K, Y.C., K.B., K.C., L.Z., I.V.N., Z.Y., M.P., S.J., D.T., and J.S.; resources: M.L., M.P., Y.D., H.C., M.P.D.M., and J.S.; writing and editing of manuscript: S.L., H.S.K., and J.S.

## Corresponding author

Correspondence to Jiaqi Shi.

---

**Supplementary Table 1.**
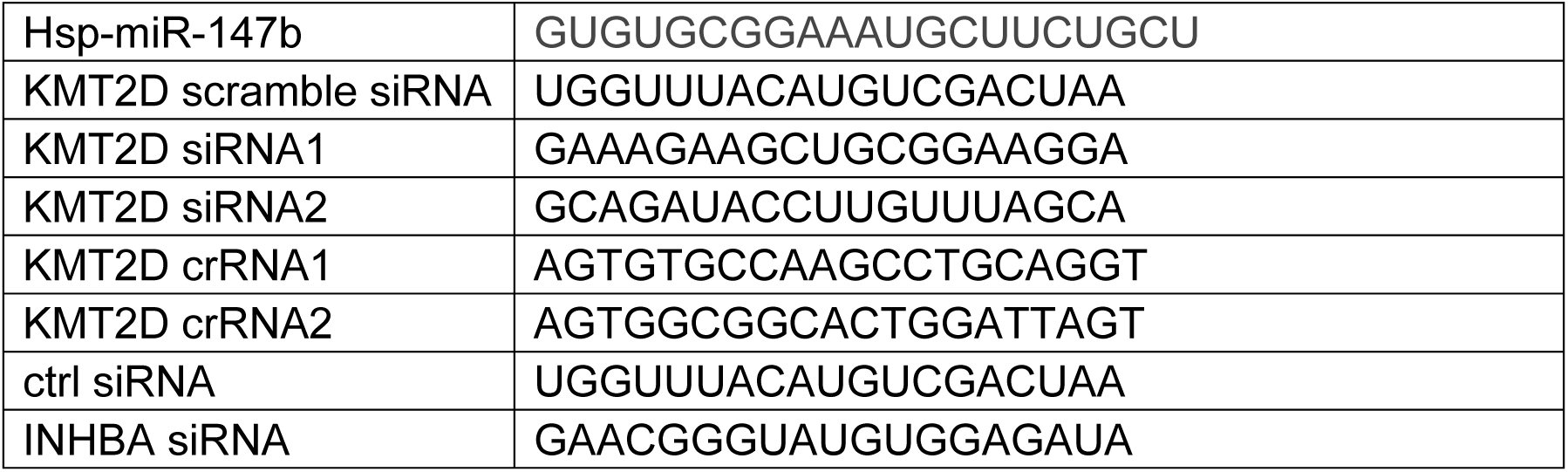
miRNA and siRNA information.

**Supplementary Table 2.**
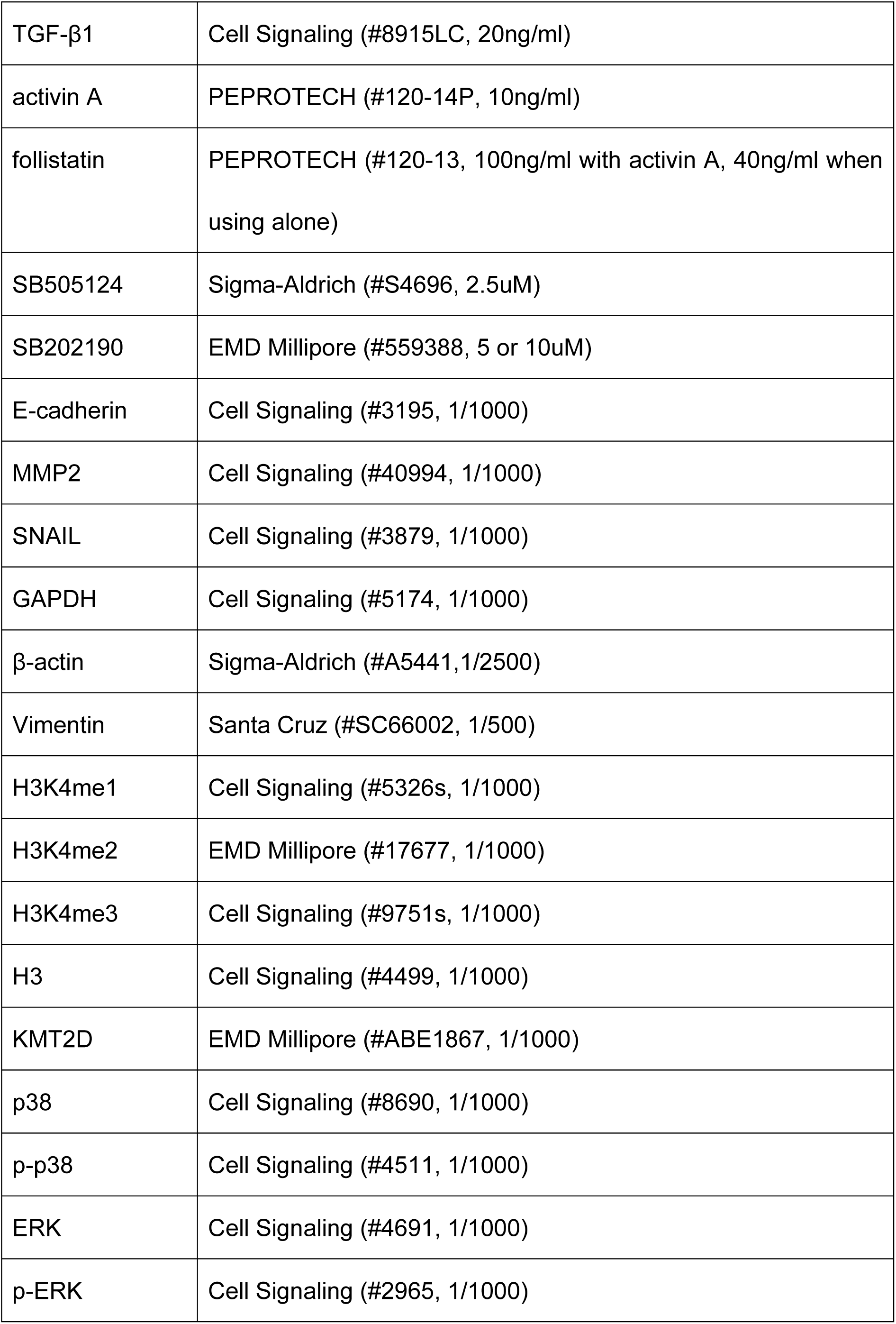
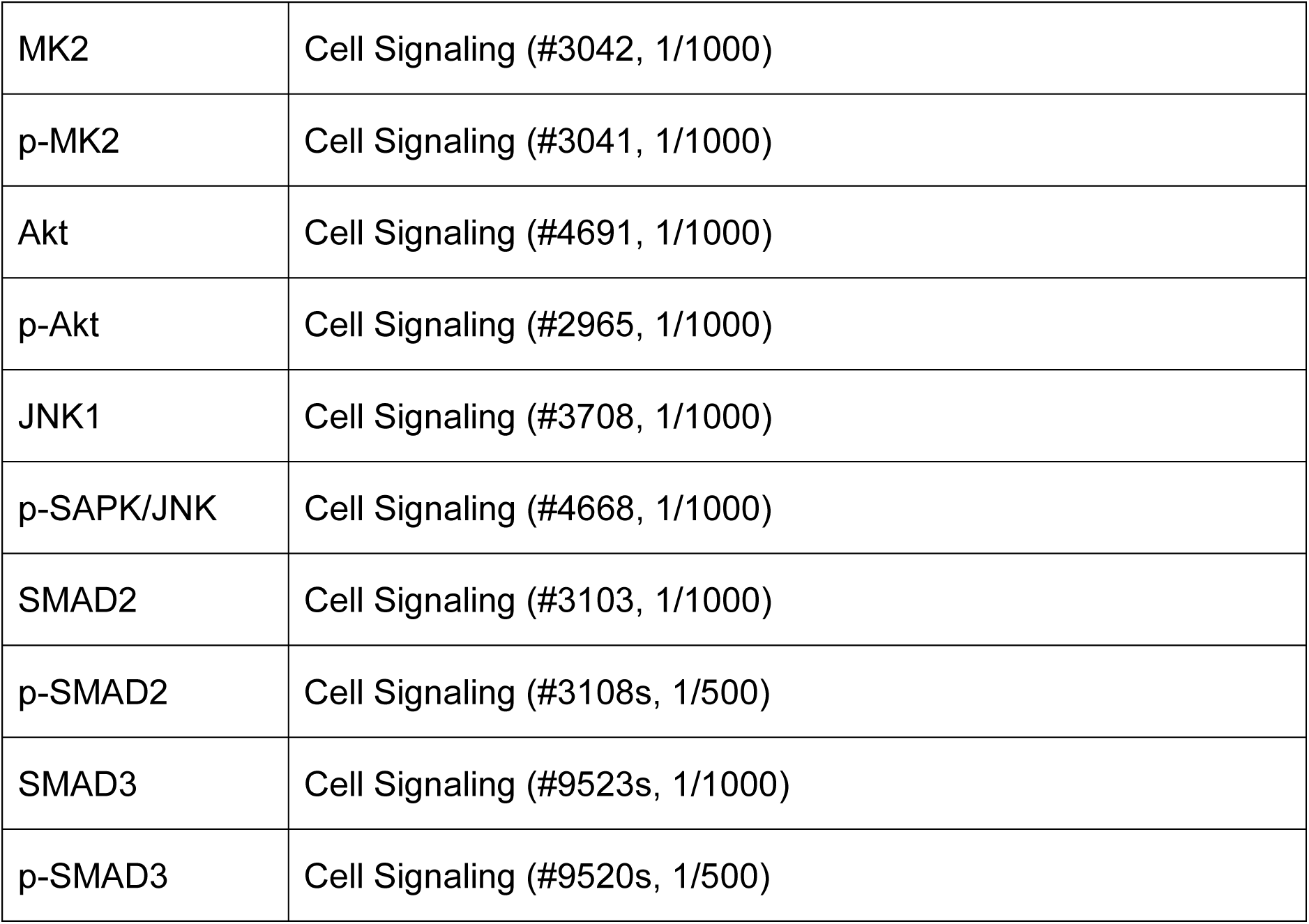
Information on antibodies and reagents.

**Supplementary Table 3.**
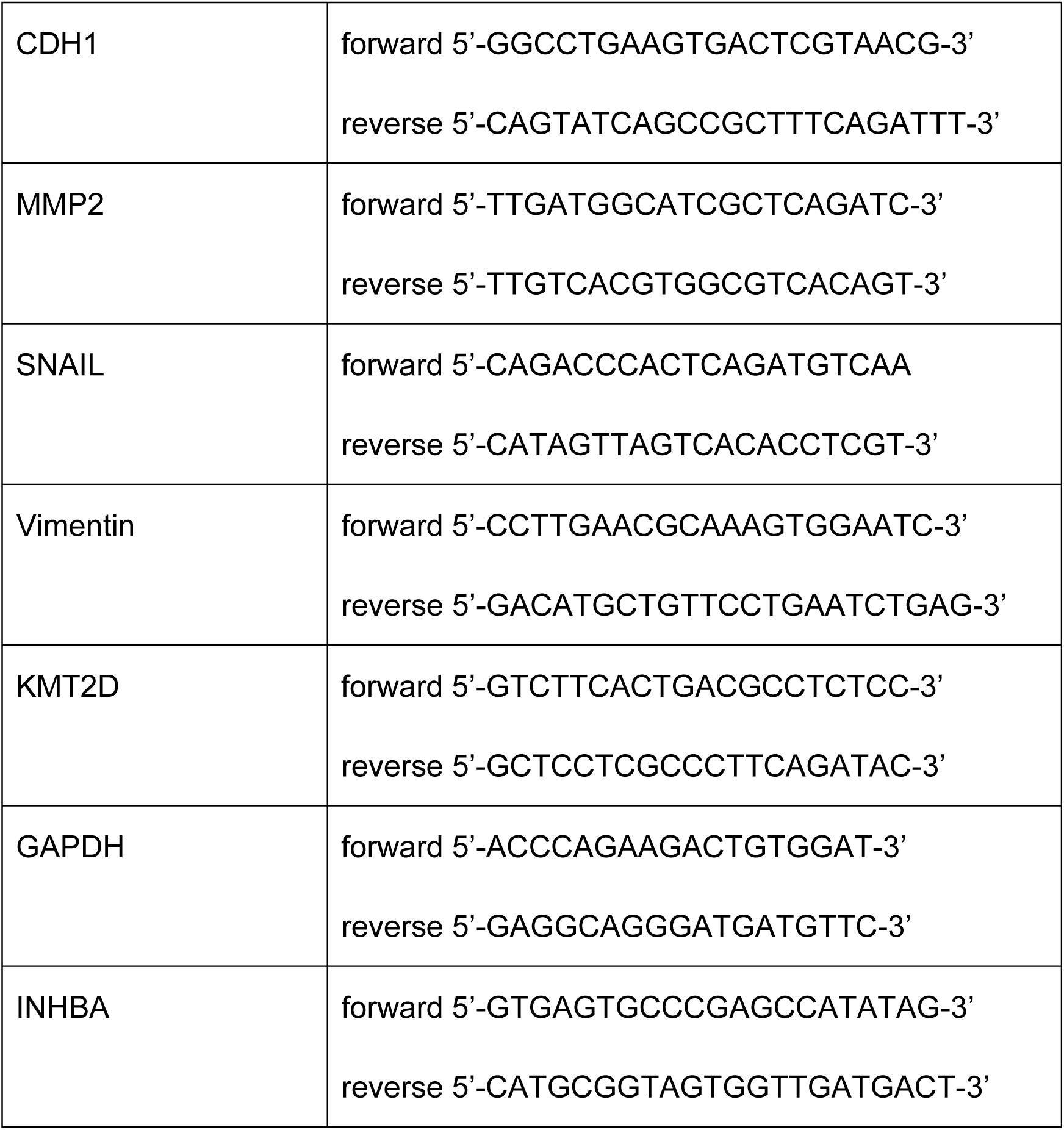
Primer sequences.

## References

1. Rahib, L. et al. Projecting cancer incidence and deaths to 2030: the unexpected burden of thyroid, liver, and pancreas cancers in the United States. Cancer Res 74, 2913–21 (2014).

2. Zhang, H. et al. MLL1 Inhibition Reprograms Epiblast Stem Cells to Naive Pluripotency. Cell Stem Cell 18, 481–94 (2016).

3. Rao, R.C. & Dou, Y. Hijacked in cancer: the KMT2 (MLL) family of methyltransferases. Nat Rev Cancer 15, 334–46 (2015).

4. Roe, J.S. et al. Enhancer Reprogramming Promotes Pancreatic Cancer Metastasis. Cell 170, 875–888 e20 (2017).

5. McDonald, O.G. et al. Epigenomic reprogramming during pancreatic cancer progression links anabolic glucose metabolism to distant metastasis. Nat Genet 49, 367–376 (2017).

6. Waddell, N. et al. Whole genomes redefine the mutational landscape of pancreatic cancer. Nature 518, 495–501 (2015).

7. Bailey, P. et al. Genomic analyses identify molecular subtypes of pancreatic cancer. Nature 531, 47–52 (2016).

8. Bird, A. DNA methylation patterns and epigenetic memory. Genes Dev 16, 6–21 (2002).

9. Zhang, J. et al. Disruption of KMT2D perturbs germinal center B cell development and promotes lymphomagenesis. Nat Med 21, 1190–8 (2015).

10. Dai, W. et al. Whole-exome sequencing reveals critical genes underlying metastasis in oesophageal squamous cell carcinoma. J Pathol 242, 500–510 (2017).

11. Yilmaz, A.S. et al. Differential mutation frequencies in metastatic cutaneous squamous cell carcinomas versus primary tumors. Cancer 123, 1184–1193 (2017).

12. Massagué, J. TGFβ signalling in context. Nature Reviews Molecular Cell Biology 13, 616 (2012).

13. Gaspar, N.J. et al. Inhibition of transforming growth factor beta signaling reduces pancreatic adenocarcinoma growth and invasiveness. Mol Pharmacol 72, 152–61 (2007).

14. Maupin, K.A. et al. Glycogene expression alterations associated with pancreatic cancer epithelial-mesenchymal transition in complementary model systems. PLoS One 5, e13002 (2010).

15. Goggins, M. et al. Genetic alterations of the transforming growth factor beta receptor genes in pancreatic and biliary adenocarcinomas. Cancer Res 58, 5329–32 (1998).

16. Hahn, S.A. et al. Homozygous deletion map at 18q21.1 in pancreatic cancer. Cancer Res 56, 490–4 (1996).

17. Hahn, S.A. et al. DPC4, a candidate tumor suppressor gene at human chromosome 18q21.1. Science 271, 350–3 (1996).

18. Heldin, C.H., Vanlandewijck, M. & Moustakas, A. Regulation of EMT by TGFbeta in cancer. FEBS Lett 586, 1959–70 (2012).

19. Paulsen, M.T. et al. Use of Bru-Seq and BruChase-Seq for genome-wide assessment of the synthesis and stability of RNA. Methods 67, 45–54 (2014).

20. Barretina, J. et al. The Cancer Cell Line Encyclopedia enables predictive modelling of anticancer drug sensitivity. Nature 483, 603–7 (2012).

21. Moffitt, R.A. et al. Virtual microdissection identifies distinct tumor- and stroma-specific subtypes of pancreatic ductal adenocarcinoma. Nat Genet 47, 1168–78 (2015).

22. Lefkofsky, H.B., Veloso, A. & Ljungman, M. Transcriptional and posttranscriptional regulation of nucleotide excision repair genes in human cells. Mutat Res 776, 9–15 (2015).

23. Loomans, H.A. & Andl, C.D. Intertwining of Activin A and TGFbeta Signaling: Dual Roles in Cancer Progression and Cancer Cell Invasion. Cancers (Basel) 7, 70–91 (2014).

24. Gurdon, J.B., Harger, P., Mitchell, A. & Lemaire, P. Activin signalling and response to a morphogen gradient. Nature 371, 487–92 (1994).

25. Hashimoto, O. et al. The role of activin type I receptors in activin A-induced growth arrest and apoptosis in mouse B-cell hybridoma cells. Cell Signal 10, 743–9 (1998).

26. Symes, K., Yordan, C. & Mercola, M. Morphological differences in Xenopus embryonic mesodermal cells are specified as an early response to distinct threshold concentrations of activin. Development 120, 2339–46 (1994).

27. McDowell, N., Zorn, A.M., Crease, D.J. & Gurdon, J.B. Activin has direct longrange signalling activity and can form a concentration gradient by diffusion. Curr Biol 7, 671–81 (1997).

28. Heldin, C.H., Landstrom, M. & Moustakas, A. Mechanism of TGF-beta signaling to growth arrest, apoptosis, and epithelial-mesenchymal transition. Curr Opin Cell Biol 21, 166–76 (2009).

29. Makohon-Moore, A.P. et al. Limited heterogeneity of known driver gene mutations among the metastases of individual patients with pancreatic cancer. Nat Genet 49, 358–366 (2017).

30. Friess, H. et al. Enhanced expression of transforming growth factor beta isoforms in pancreatic cancer correlates with decreased survival. Gastroenterology 105, 1846–56 (1993).

31. David, C.J. et al. TGF-beta Tumor Suppression through a Lethal EMT. Cell 164, 1015–30 (2016).

32. Li, S., Wu, L., Wang, Q., Li, Y. & Wang, X. KDM4B promotes epithelialmesenchymal transition through up-regulation of ZEB1 in pancreatic cancer. Acta Biochim Biophys Sin (Shanghai) 47, 997–1004 (2015).

33. Ramadoss, S., Chen, X. & Wang, C.Y. Histone demethylase KDM6B promotes epithelial-mesenchymal transition. J Biol Chem 287, 44508–17 (2012).

34. Volinia, S. et al. A microRNA expression signature of human solid tumors defines cancer gene targets. Proc Natl Acad Sci U S A 103, 2257–61 (2006).

35. Roldo, C. et al. MicroRNA expression abnormalities in pancreatic endocrine and acinar tumors are associated with distinctive pathologic features and clinical behavior. J Clin Oncol 24, 4677–84 (2006).

36. Bloomston, M. et al. MicroRNA expression patterns to differentiate pancreatic adenocarcinoma from normal pancreas and chronic pancreatitis. Jama 297, 1901–8 (2007).

37. Zhang, E. et al. MicroRNA miR-147b promotes tumor growth via targeting UBE2N in hepatocellular carcinoma. Oncotarget 8, 114072–114080 (2017).

38. Augello, C. et al. MicroRNA as potential biomarker in HCV-associated diffuse large B-cell lymphoma. J Clin Pathol 67, 697–701 (2014).

39. Zhang, W.C. et al. miR-147b-mediated TCA cycle dysfunction and pseudohypoxia initiate drug tolerance to EGFR inhibitors in lung adenocarcinoma. Nat Metab 1, 460–474 (2019).

40. Ford, D.J. & Dingwall, A.K. The cancer COMPASS: navigating the functions of MLL complexes in cancer. Cancer Genet 208, 178–91 (2015).

41. Ardeshir-Larijani, F. et al. KMT2D Mutation Is Associated With Poor Prognosis in Non-Small-Cell Lung Cancer. Clin Lung Cancer 19, e489–e501 (2018).

42. Abudureheman, A. et al. High MLL2 expression predicts poor prognosis and promotes tumor progression by inducing EMT in esophageal squamous cell carcinoma. J Cancer Res Clin Oncol 144, 1025–1035 (2018).

43. Dawkins, J.B. et al. Reduced Expression of Histone Methyltransferases KMT2C and KMT2D Correlates with Improved Outcome in Pancreatic Ductal Adenocarcinoma. Cancer Res 76, 4861–71 (2016).

44. Koutsioumpa, M. et al. Lysine methyltransferase 2D regulates pancreatic carcinogenesis through metabolic reprogramming. Gut 68, 1271–1286 (2019).

45. Camolotto, S.A., Belova, V.K. & Snyder, E.L. The role of lineage specifiers in pancreatic ductal adenocarcinoma. J Gastrointest Oncol 9, 1005–1013 (2018).

46. Juiz, N.A., Iovanna, J. & Dusetti, N. Pancreatic Cancer Heterogeneity Can Be Explained Beyond the Genome. Front Oncol 9, 246 (2019).

47. Smith, J.C., Price, B.M., Van Nimmen, K. & Huylebroeck, D. Identification of a potent Xenopus mesoderm-inducing factor as a homologue of activin A. Nature 345, 729–31 (1990).

48. Nazareth, E.J.P., Rahman, N., Yin, T. & Zandstra, P.W. A Multi-Lineage Screen Reveals mTORC1 Inhibition Enhances Human Pluripotent Stem Cell Mesendoderm and Blood Progenitor Production. Stem Cell Reports 6, 679–691 (2016).

49. Green, J.B., New, H.V. & Smith, J.C. Responses of embryonic Xenopus cells to activin and FGF are separated by multiple dose thresholds and correspond to distinct axes of the mesoderm. Cell 71, 731–9 (1992).

50. Katz, L.H. et al. Targeting TGF-beta signaling in cancer. Expert Opin Ther Targets 17, 743–60 (2013).

51. Paulsen, M.T. et al. Coordinated regulation of synthesis and stability of RNA during the acute TNF-induced proinflammatory response. Proc Natl Acad Sci U S A 110, 2240–5 (2013).

52. Bedi, K., Paulsen, M.T., Wilson, T.E. & Ljungman, M. Characterization of novel primary miRNA transcription units in human cells using Bru-seq nascent RNA sequencing. NAR Genom Bioinform 2, lqz014 (2020).

53. Bereshchenko, O.R., Gu, W. & Dalla-Favera, R. Acetylation inactivates the transcriptional repressor BCL6. Nat Genet 32, 606–13 (2002).

54. Nguyen, N. et al. Tumor infiltrating lymphocytes and survival in patients with head and neck squamous cell carcinoma. Head Neck 38, 1074–84 (2016).

