## Supplementary Information for "KMT2D links TGF-β Signalling to Non-Canonical Activin Pathway and Regulates Pancreatic Cancer Cell Plasticity"

**Lu et al.**

Supplementary Information

Supplementary Figure 1

Supplementary Figure 2

Supplementary Figure 3

Supplementary Figure 4

### Supplementary Figure 1

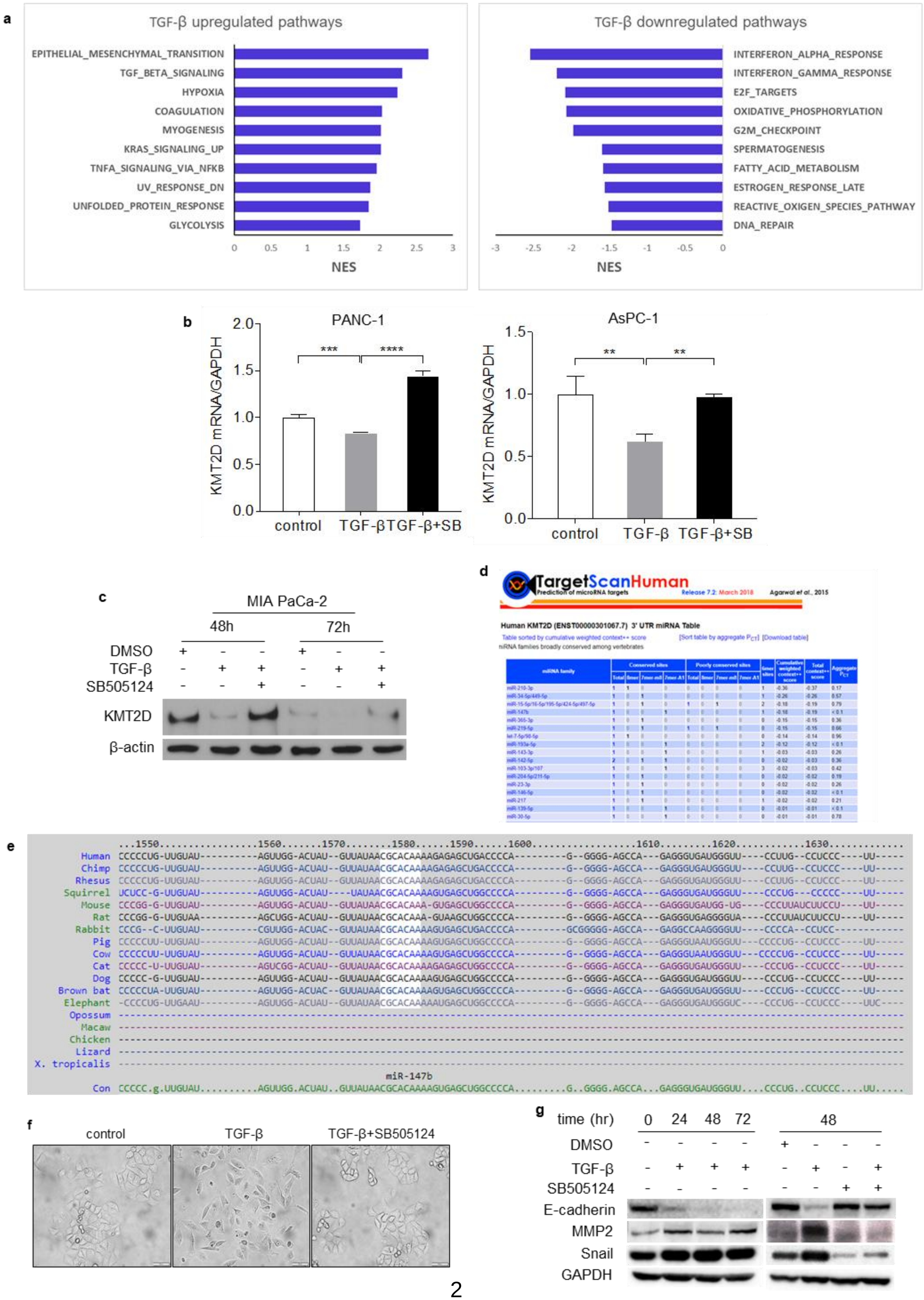

**Supplementary Fig. 1** TGF-β downregulates KMT2D and promotes EMT in PDAC cells. **a** Top hallmark pathways that are up- or down-regulated in 72h TGF-β treated PANC-1 cells. NES: normalized enrichment score. **b** mRNA expression of KMT2D in PANC-1 and AsPC-1 cells treated with TGF-β with or without its inhibitor SB505124. (**$p<0.01$, ***$p<0.005$, ****$p<0.001$, One-way ANOVA test with Dunnett's multiple comparisons test, n=3) **c** Western Blot analysis of MIA PaCa-2 cells treated with 20ng/ml TGF-β with or without 2.5nM SB505124 or DMSO for 48h or 72h. β-actin was used as loading control. **d** TargetScan prediction of potential miRNAs targeting 3' UTR of KMT2D. **e** Conserved binding site of miR-147b among mammals queried from TargetScanHuman. **f** Phase-contrast images of PANC-1 cells treated with DMSO, or TGF-β with or without SB505124. (scale bar = 100 um). **g** Western Blot analysis of E-cadherin, MMP2, and Snail in PANC-1 cells treated with DMSO or 20 ng/ml TGF-β with or without SB505124 for 24, 48, and 72 hours. GAPDH was used as loading control.

### Supplementary Figure 2

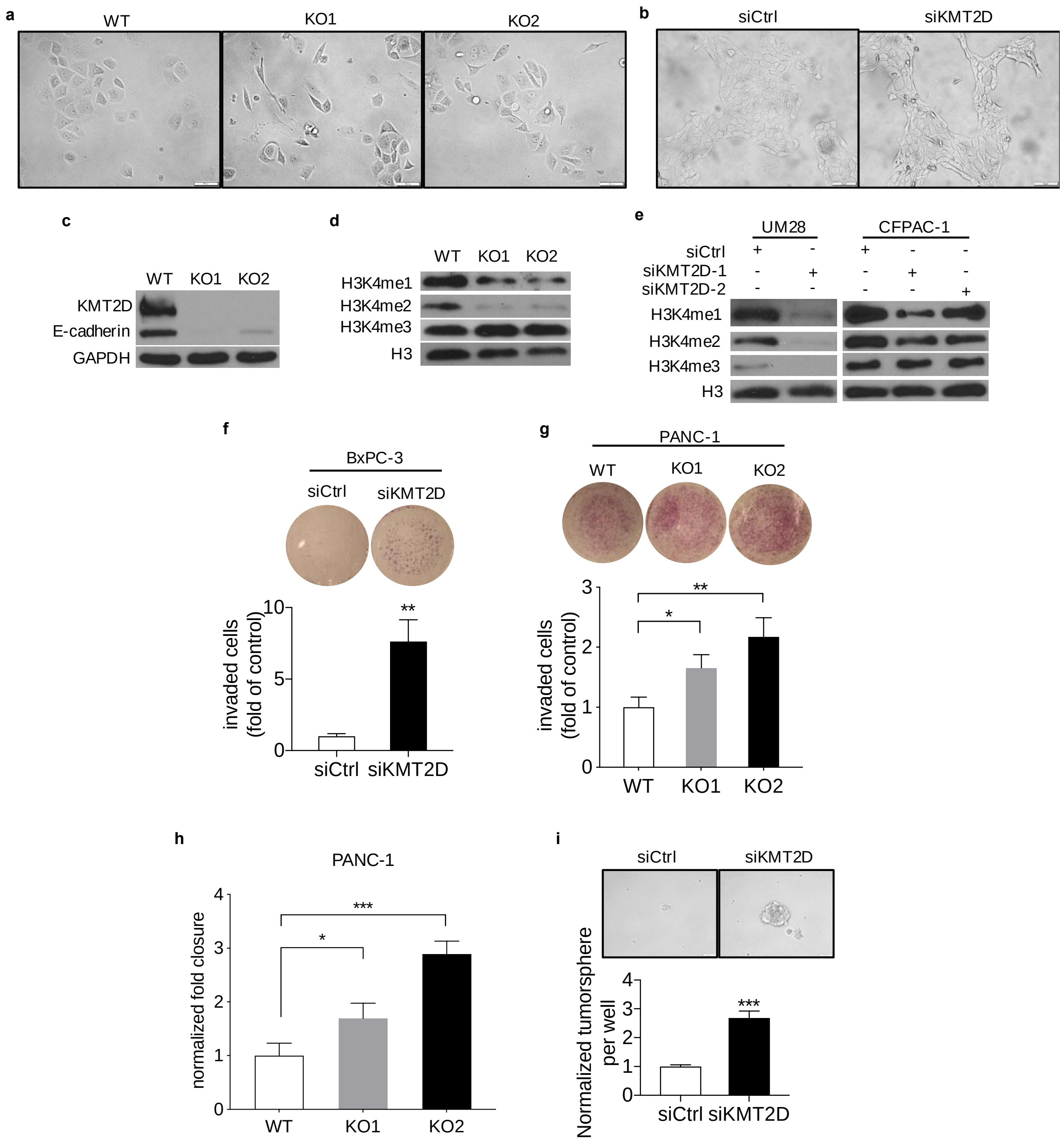


**Supplementary Fig. 2** Reduced KMT2D expression promotes EMT, migration, invasion, and tumorigenicity in PDAC cells. **a** Phase-contrast images of PANC-1 wild-type (WT) cells and KMT2D knockout (KO) cells. Scale bar: 100um. **b** Phase-contrast images of BxPC-3 cells transfected with 50nM scramble siRNA (siCtrl) or KMT2D siRNA (siKMT2D) for 5 days. **c** Western Blot analysis of KMT2D and E-cadherin in KMT2D KO PANC-1 cells compared to WT PANC-1 cells. GAPDH was used as loading control. **d** Western blot of histone H3 lysine4 methylation levels in WT and KMT2D KO BxPC-3 cells. H3 was used as loading control. **e** Western Blot analysis of histone H3 lysine4 methylation levels in siCtrl or siKMT2D transfected UM28 or CFPAC-1 PDAC cell lines. H3 was used as loading control. **f** Invaded BxPC-3 cells transfected with either siCtrl or siKMT2D at 48h by transwell invasion assay (**$p<0.01$, unpaired student t-test, n=3). **g** Invaded cells in 2 KMT2D KO PANC-1 clones and WT PANC-1 cells at 48 h by transwell invasion assay. (*$p<0.05$, **$p<0.01$, one-way ANOVA with Dunnett's multiple comparisons test, n=3) **h** Wound closure in KMT2D KO PANC-1 cells and WT PANC-1 cells at 48h post scratch by wound healing. **i** Tumorspheres formed by siCtrl or siKMT2D transfected UM28 cells by tumorsphere formation assay. (***$p<0.005$, unpaired student t-test, n=3)

### Supplementary Figure 3

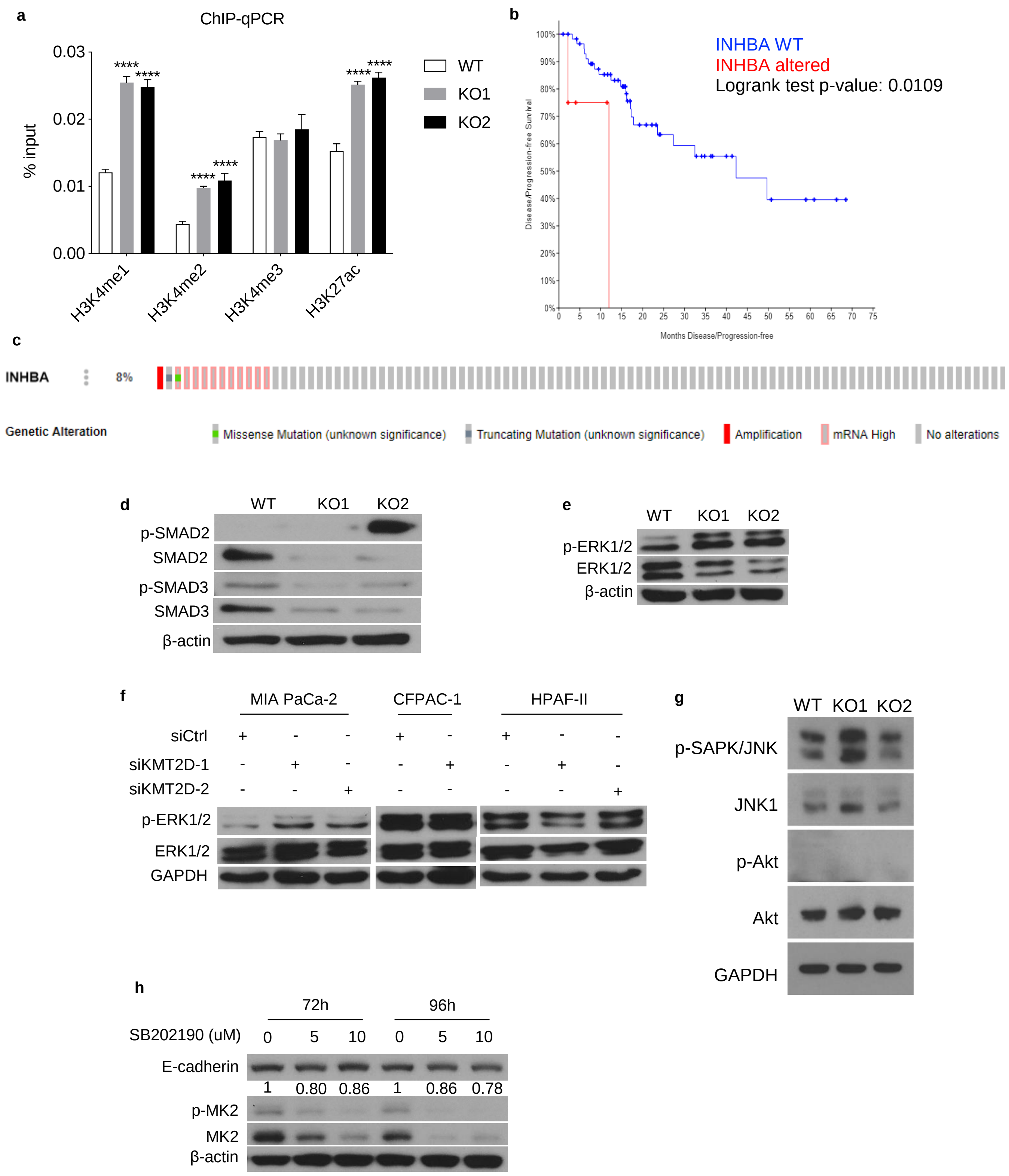


**Supplementary Fig. 3** KMT2D depletion activates the non-canonical activin A pathway. **a** Chromatin immunoprecipitation (ChIP) and real-time PCR analysis of histone modifications at the INHBA promoter of WT and KMT2D KO BxPC-3 cells. Results were normalized to inputs and expressed as % input. **b** Kaplan-Meir curve of disease-free survival in PDAC patients with or without alterations in INHBA (n=168, TCGA database). **c** Genetic alterations of INHBA in PDAC tissues (n=168, TCGA database). **d** Western blot analysis of p-SMAD2, SMAD2, p-SMAD3, and SMAD3 in KMT2D WT and KO BxPC-3 cells. B-actin was used as loading control. **e** Western Blot analysis of ERK1/2 and p-ERK1/2 in WT and KMT2D KO BxPC-3 cells. β-actin was used as loading control. **f** Western Blot analysis of ERK and p-ERK1/2 in PDAC cells transfected with siCtrl or siKMT2D. GAPDH was used as loading control. **g** Western blot analysis of p-SAPK/JNK, JNK, p-Akt, and Akt in WT and KMT2D KO BxPC-3 cells. GAPDH was used as loading control. **h** Western Blot analysis of E-cadherin, p-MK2, and MK2 in wild-type BxPC-3 cells treated with either 5 or 10 uM SB202190 or DMSO control for 72h or 96h. B-actin was used as loading control.

### Supplementary Figure 4

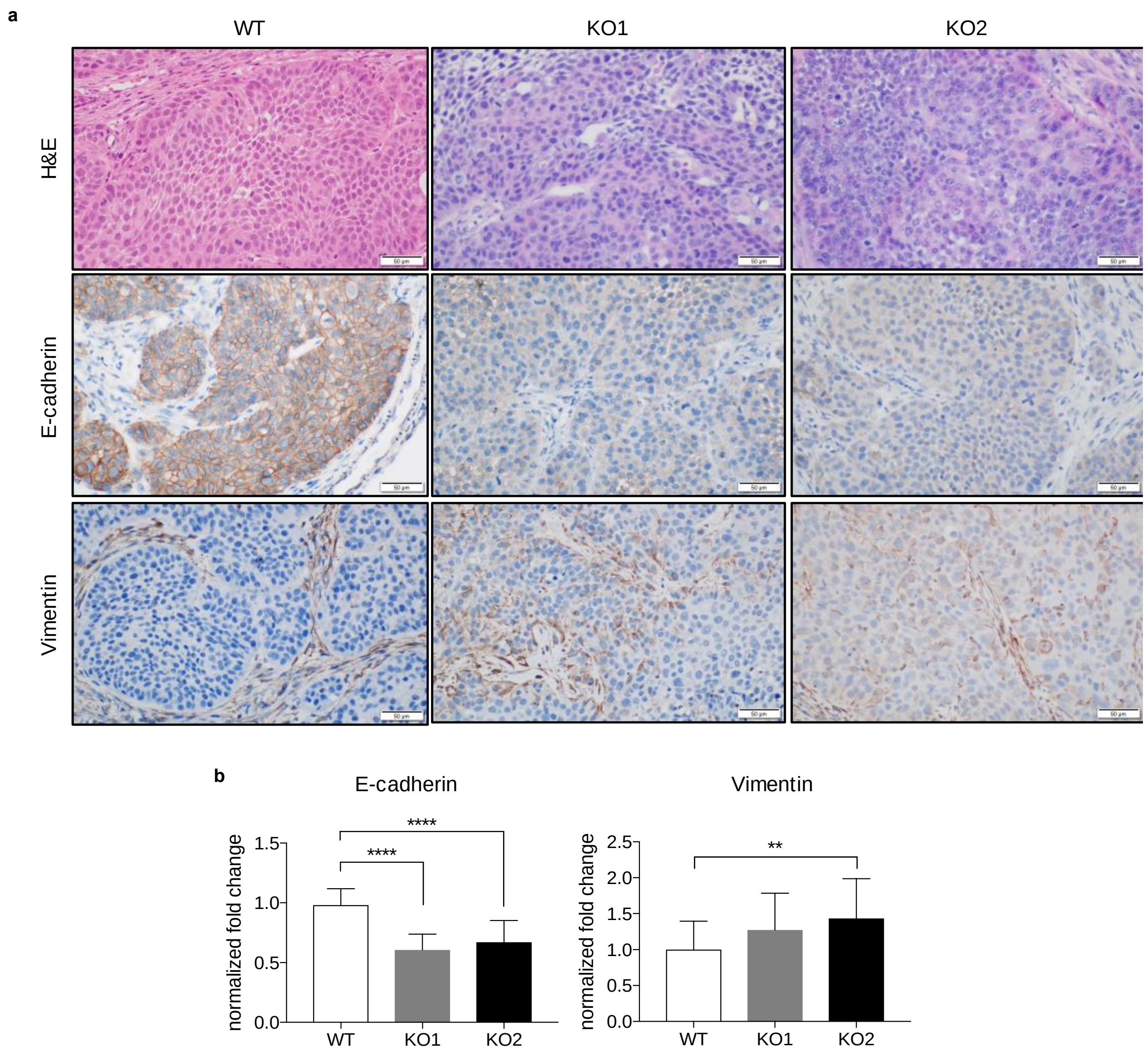


**Supplementary Fig. 4** KMT2D knockout cells form tumors with enhanced mesenchymal morphology in vivo. **a** Representative histology and immunohistochemical stains of E-cadherin and Vimentin in BxPC-3 WT and KMT2D-KO orthotopic xenograft tumors. (scale bar = 50 um) **b** Quantification of immunohistochemistry stains of E-cadherin and Vimentin in WT and KMT2D KO xenograft tumors shown in **a**. (****p<0.001, One-way ANOVA test with Tukey's multiple comparisons, n=10)
